# *NF2* loss malignantly transforms human pancreatic acinar cells and enhances cell fitness under environmental stress

**DOI:** 10.1101/2025.04.29.651251

**Authors:** Yi Xu, Michael H. Nipper, Angel A. Dominguez, Chenhui He, Francis E. Sharkey, Sajid Khan, Han Xu, Daohong Zhou, Lei Zheng, Yu Luan, Jun Liu, Pei Wang

## Abstract

Pancreatic ductal adenocarcinoma (PDAC) occurs as a complex, multifaceted event driven by the interplay of tumor permissive genetic mutations, nature of cellular origin and microenvironmental stress. In this study, using primary human pancreatic acinar 3D organoids, we performed CRISPR knockout screen targeting 199 previously underappreciated potential tumor suppressors curated from clinical PDAC samples. Our data revealed significant enrichment of a list of candidates, with NF2 emerging as the top target. Functional validation confirmed that loss of NF2 promotes the transition of PDAC to an invasive state, potentially through extracellular matrix modulation. NF2 inactivation was found to enhance PDAC cell fitness under nutrient starvation. This adaptation not only reinforces the oncogenic state but also confers therapeutical resistance. Additionally, we found that NF2 loss is associated with the fibroblast heterogeneity and cancer-stroma communications in tumor evolution. These findings establish *NF2* as a critical tumor suppressor in PDAC and uncover its role in mediating nutrient adaptation and drug resistance. Importantly, this study provides new insights into drug resistance mechanisms and potential therapeutic targets in PDAC.

## INTRODUCTION

Pancreatic Ductal Adenocarcinoma (PDAC) remains one of the most lethal malignancies, with a dismal 5-year survival rate of approximately 13% (1). Many contributing factors including genetic mutations, the nature of cell of origin and tumor microenvironmental signaling drive PDAC tumorigenesis (2–4). Over 90% of PDAC patients harbor oncogenic mutations in *KRAS*, with additional inactivating mutations in tumor suppressor genes including *CDKN2A*, *TP53* or *SMAD4* commonly observed in PDAC patients. These oncogenic alterations provide a selective growth advantage that facilitates tumor initiation and progression, and as such, are classified as PDAC driver mutations. Despite recent advances in targeted therapies aimed at oncogenic *RAS* signaling (5, 6), therapeutic strategies targeting other key driver mutations remain limited.

Many PDAC patients do not carry all four key driver mutations simultaneously, suggesting the presence of additional oncogenic events contributing to disease progression. Indeed, next-generation sequencing has identified thousands of somatic mutations in PDAC, with over one hundred mutated genes on average identified in each patient (7–13). While most of these mutations may represent passenger mutations which do not contribute to PDAC pathogenesis, it is very likely that some previously underappreciated recurrent mutations in PDAC patients also function as cancer driver mutations. Identifying these potentially new PDAC driver genes is a key step toward understanding tumor biology and developing targeted therapies. In addition, establishing a system that is efficient and cost effective for identification of the causal relationship between gene mutations and therapeutic resistance will offer potential targets to improve treatment outcome of human PDAC.

We have previously reported a flow cytometry-based method to isolate primary acinar and ductal cells from healthy human pancreas tissues, which can be engineered to recapitulate human PDAC early development (14–17). Using engineered human cells as a cancer model has inherent advantages over mouse models and established cancer cell lines in the investigation of disease initiation and early progression, thus providing an effective platform to identify novel PDAC drivers in a lineage specific manner. In the present study, we combined our unique early human PDAC model with CRISPR screening to perform a knockout screen targeting 199 potential tumor suppressor driver mutations in engineered human primary pancreatic acinar cells. Analysis of the screen results identified *NF2* loss as a prominent driver mutation that promotes acinar-derived PDAC progression. Further validation and mechanistic studies revealed that *NF2* inactivation induces intrinsic transcriptomic changes that can prime acinar cells to enhance their fitness under nutrient deprivation, ultimately leading to multidrug resistance.

## RESULTS

### 1. Unveil potential cancer driver mutations in tumor suppressors using primary human pancreatic acinar cells

We employed our previously established protocol to isolate primary acinar cells from healthy human pancreatic tissues (14, 15) (donor information in **Supplementary Table 1**), which were maintained as 3D organoids for long-term culture (**Figure 1A**). To accelerate acinar transformation for proposed screen, the organoids were engineered to carry three key PDAC driver mutations: overexpression of oncogenic *KRAS^G12V^*, and knockout of *TP53* and *CDKN2A* (designated as KPT organoids, **Figure S1A-B**). This combination was determined to be the threshold mutation burden required for acinar transformation in our model (15). Through a comprehensive literature search, we curated a list of 199 potential tumor suppressors recurrently inactivated in clinical PDAC samples (7–13). A pooled CRISPR knockout sgRNA library targeting these genes was constructed (**Supplementary Table 2**) and validated (**Figure S1C**). The library was then packaged into lentivirus and introduced into four independently established KPT organoid cultures with duplicates for each (**Figure 1B**). Following antibiotic selection and a brief expansion, the organoids were transplanted into NSG immunodeficient mice (NOD/SCID/gamma, NOD.Cg-Prkdcscid IL2Rγtm1Wjl/SzJ, JAX Lab) for 8 weeks. Tumors were subsequently collected and subjected to next-generation sequencing (NGS).

**Figure 1.**
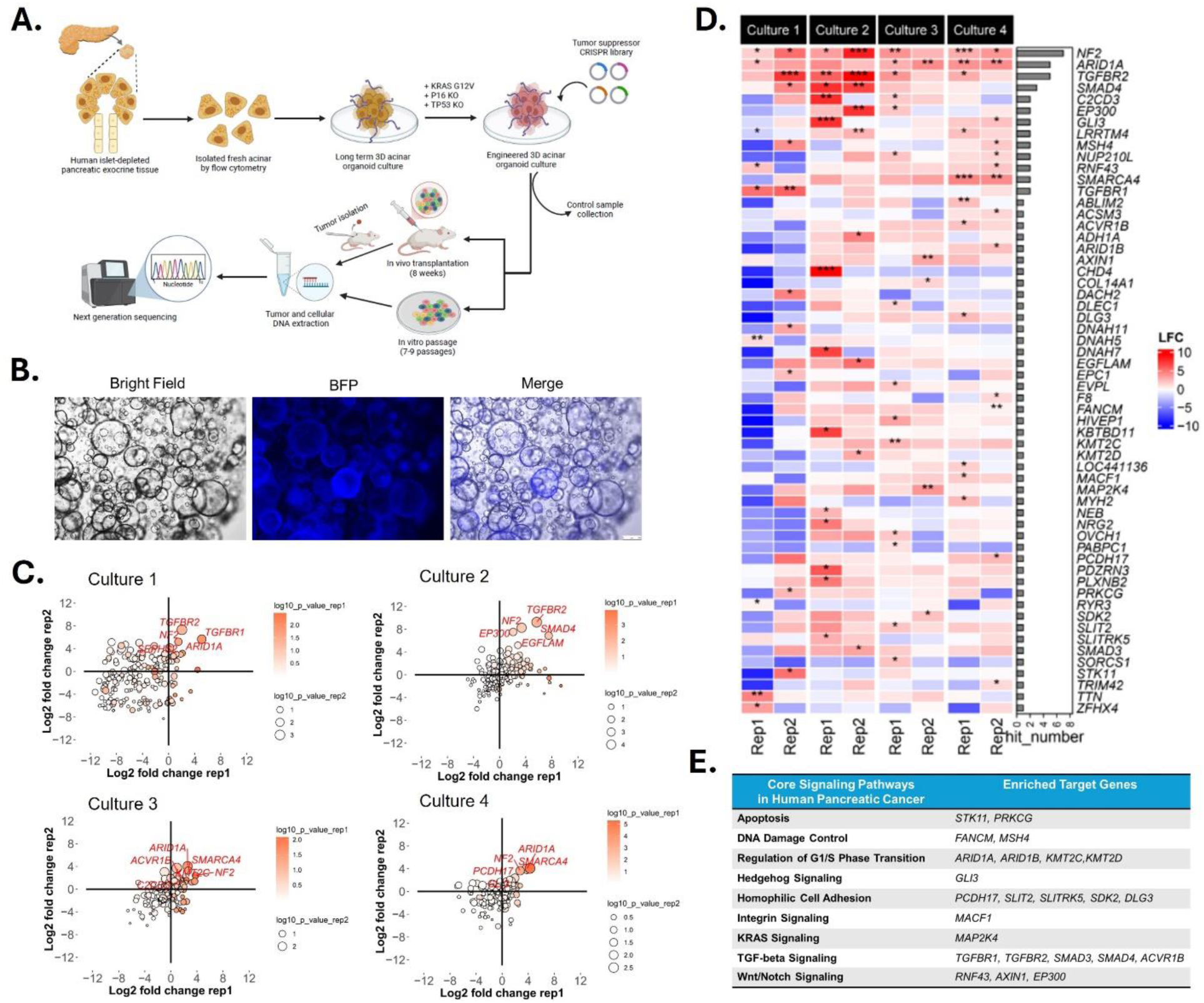
Unveil potential tumor suppressor driver mutations in human PDAC using primary pancreatic acinar cells. **A.** Schematic illustration of CRISPR knockout screen using human primary acinar 3D organoid culture. **B.** Bright field and fluorescence images of acinar organoids transduced with lentivirus expressing BFP tagged CRISPR sgRNA library, scale bar = 250 µm. **C.** Scatter plot of the enrichment of target genes from *in vivo* CRISPR screen in 4 independent cultures with 2 replicates each. X and Y axis represent log2 fold change of the sgRNAs distribution of an indicated target gene in individual replicate compared with the control. **D.** Heatmap of all positively enriched target genes in any of the 4 independent cultures. The color scale represents log2 fold change (LFC). The side bar plot indicates the number of replicates in which each target was found to be enriched. *, **, *** indicate p-value < 0.05, 0.01, 0.001, respectively. **E.** The core signaling pathways in human pancreatic cancer in which indicated target genes are involved.

Analysis of NGS data revealed significant enrichment of sgRNAs targeting 58 genes in at least one replicate, suggesting potential advantages in tumor progression upon loss of these target genes (**Figure 1C-D, Supplementary Table 3**). Notably, *SMAD4*, an established PDAC driver that was included in the library as a positive control, was found to be one of the top enriched targets. On the other hand, another positive control target *TP53* was not enriched in any of the replicates as *TP53* inactivation was already introduced in the KPT organoids as our starting materials. These observations together demonstrated the robustness of our methodology. Furthermore, 24 out of the 58 genes from the list could be functionally assigned to one of the genetically altered core signaling pathways identified by whole genome sequencing in human pancreatic cancer patients (7), suggesting their contributions in PDAC development (**Figure 1E**). In comparison, the implications of the other 34 genes in PDAC are less understood.

Specifically, among all the enriched targets, *NF2*, *ARID1A* and *TGFBR2* emerged as the top 3 enriched targets which were identified in more than half of the replicates, highlighting their significant impact on tumor development. *ARID1A* has been previously reported to maintain acinar homeostasis and prevent PDAC transformation in engineered mouse models (18), while loss of *TGFBR2* contributes to tumor development mainly through the TGFβ signaling pathway. In comparison, although the tumor suppressive role of *NF2* has been established in mesothelioma, schwannomas and meningiomas where it is frequently mutated(19, 20), the contributions of *NF2* inactivation to PDAC development remain elusive.

In addition, we also performed an *in vitro* screening to assess the influence of environmental context on screen outcomes. Transduced organoids were maintained *in vitro* for 8 weeks (∼ 7-9 passages), followed by DNA extraction and NGS analysis to compare their library distributions with those of freshly transduced cells. This analysis revealed significant enrichment of sgRNAs targeting 31 genes in at least one replicate (**Figure S1D-E, Supplementary Table 3**). As anticipated, there was a considerable disparity between the enriched target genes identified from the *in vitro* versus *in vivo* screen (**Figure S1F**), suggesting a potential interplay between environmental context and genetic predisposition during tumor development. Nevertheless, a total of 13 genes were identified in both settings, with *NF2* remaining as the top hit in both conditions, indicating a strong growth advantage conferred by *NF2* loss in transformed acinar cells.

### 2. Loss of *NF2* facilitates aggressive progression of acinar-derived human PDAC

*NF2* encodes merlin, a cytoskeletal protein that regulates cell-cell contact inhibition and cellular interactions with extracellular matrix (ECM) signaling. Analysis of the TCGA PDAC cohort revealed that lower *NF2* expression correlates with a poorer prognosis, especially in the patients associated with a classical subtype (21), suggesting a tumor suppressive role from this gene (**Figure 2A**). To investigate the contribution of *NF2* inactivation to the development of acinar-derived PDAC, we generated *NF2*-knockout acinar cultures (designated as KPTN) in four independently established KPT organoid cultures which were verified by Western blot at the protein level (**Figure 2B**). When the KPTN cells were co-cultured with KPT cells at a 1:1 ratio *in vitro*, KPTN cells gradually outcompeted KPT cells and reached dominance by passage 7 (**Figure S2A**). This observation aligns with the growth advantage conferred by *NF2* knockout observed in *in vitro* CRISPR screen.

**Figure 2.**
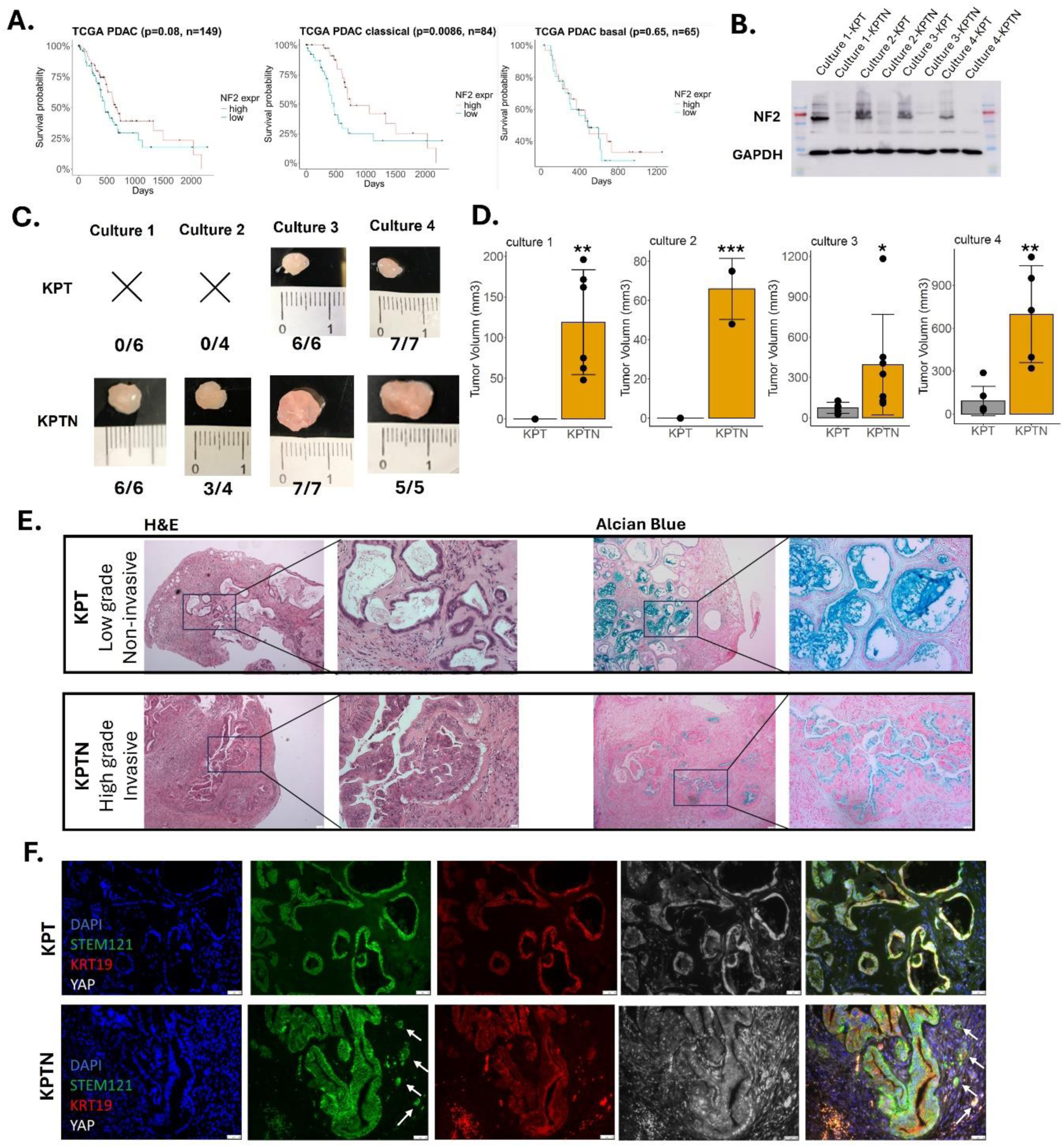
Loss of *NF2* facilitates aggressive progression of acinar-derived human PDAC. **A.** Kaplan-Meier plots of TCGA PDAC patients (all 149 patients, 84 classical PDAC patients, 65 basal PDAC patients, respectively) separated by median *NF2* expression level. **B.** Western blot of *NF2* expression in KPT and KPTN organoids derived from 4 independent cultures. **C.** Representative images of xenograft tumors harvested from NSG mice transplanted with KPT and KPTN organoids derived from 4 independent cultures. The number of tumor formation/ transplantations was indicated in the figure. **D.** Quantification of tumor size. Data was analyzed by two-tailed Student’s t test. Error bar represents standard deviation. *, **, *** indicate p-value < 0.05, 0.01, 0.001, respectively, compared with corresponding KPT counterparts. **E.** Representative HE and Alcian blue staining of tumor sections. Scale bar = 100 µm. Insert scale bar = 25 µm. **F.** Representative immunofluorescence staining of tumor sections. Cell nuclei was counter stained with DAPI. Scale bar = 50 µm. Arrows indicate invasive lesions.

To assess the impact of *NF2* inactivation on tumorigenicity, the engineered organoids were subcutaneously transplanted into NSG mice. KPT cells derived from cultures 1 (n = 6) and 2 (n = 4) failed to form tumors within 8 weeks of inoculation. In contrast, *NF2* knockout in these cultures resulted in successful tumor formation (n = 6 and 4, respectively) (**Figure 2C**, **S2B**). Notably, while xenograft tumors were successfully established by transplanting KPT cells from cultures 3 (n = 6) and 4 (n = 7), *NF2* inactivation significantly increased the tumor size compared to their KPT counterparts (n = 7 and 5, respectively) (**Figure 2C-D**, **S2B**). Histological analysis of tumor tissue sections revealed that KPTN tumors progressed to high grade lesions characterized by nuclear atypia, whereas all KPT tumors remained low grade lesions with substantial Alcian blue staining (**Figure 2E**). Immunofluorescence staining with a human specific STEM121 antibody marked the tumor cells of human origin and further confirmed the presence of invasive cancer cell populations in KPTN tumors (**Figure 2F, S2C**). These findings together suggested that loss of *NF2* facilitated tumor development and promoted malignant progression of acinar-derived PDAC. Consistent with our previous findings (15), all the acinar-derived tumor cells were associated with strong expression of KRT19, a marker of normal ductal lineage not expressed in healthy acinar cells. In addition, in line with the established role of *NF2* in the hippo pathway, KPTN tumors displayed abundant nuclear YAP1 staining, further validating the functional consequences of *NF2* knockout.

### 3. Molecular dissecting the *NF2* loss-induced tumor evolution under *in vivo* context

To understand how *NF2* inactivation can promote acinar-derived PDAC transformation, bulk RNA-seq was performed to detect global transcriptional alterations in KPTN versus KPT-derived tumor tissues (n = 3). We identified 683 human genes significantly upregulated in KPTN tumors (fold change > 2, adjusted P value < 0.05), which are involved in proliferation, glycolysis, hypoxia, focal adhesion, as well as collagen formation, suggesting a more active cancer progression (**Figure 3A-B, Supplementary Table 4**). Conversely, 476 genes downregulated in KPTN tumors included immune related genes, consistent with our previous report (15), highlighting a possible intrinsic immune evasion mechanism during PDAC progression (**Figure 3B**).

**Figure 3.**
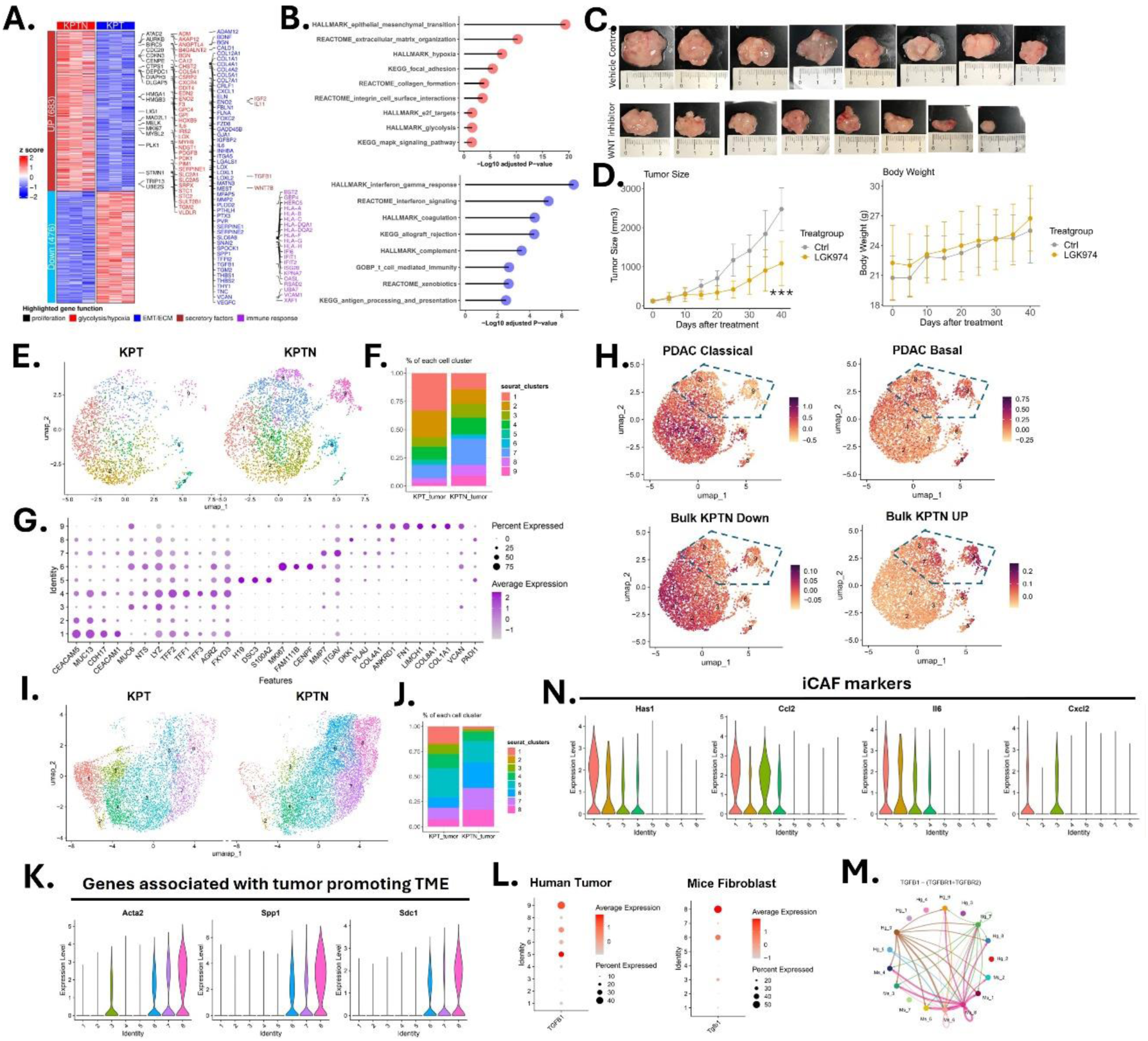
Molecular dissecting the *NF2* loss-induced tumor evolution under in vivo context. **A.** Expression heatmap of differentially expressed genes in KPTN vs KPT tumors (n =3 in each group, fold change > 2, adj.p < 0.05). Side annotation included genes involved in specific functional pathways as indicated at the bottom. **B.** Overrepresentation analysis for upregulated (in red) and downregulated genes (in blue) in KPTN tumor as shown in **A**. A significant enrichment at pathway level was considered with multiple-test adjusted p-value < 0.05. **C.** Photos of xenograft tumors from NSG mice transplanted with KPTN tumors and received LGK974 at a daily dose of 10 mg/kg (oral gavage) or vehicle control. **D.** Tumor size and body weight comparison between mice received LGK974 or vehicle control. *** indicate p-value < 0.001 compared with vehicle treatment group. **E.** UMAP of cells of human origin present in KPT and KPTN-derived tumor tissues in the single cell RNA seq analysis. **F.** Percentage composition of each cell population as shown in **E**. **G.** Expressions of indicated feature genes in each cell population as shown in **E**. **H.** Expression level of indicated gene modules in all the populations as shown in **E**. **I.** UMAP of all mice fibroblast cells present in KPT and KPTN-derived tumor tissues. **J.** Percentage composition of each cell population as shown in **I**. **K.** Violin plot of the indicated gene expression associated with tumor promoting TME in each cell population as shown in **I**. **L.** *TGFB1* and *tgfb1* gene expression in each cell populations of human and mice origin, respectively. **M.** Inferred cell-cell communication via TGFB1 signaling among cell populations shown in **I**. **N.** Violin plot of the indicated gene expression associated with iCAF in each cell population as shown in **I**.

The bulk RNA-seq data revealed *WNT7B* upregulation in KPTN tumors, an essential niche factor for maintaining human pancreatic organoid culture as well as supporting tumor growth (22, 23). To test its therapeutic potential, we treated KPTN xenografts in NSG mice with the WNT secretion inhibitor LGK974 (10 mg/kg daily) (24). After 40 days, LGK974 significantly reduced tumor size compared with the vehicle treatment group, without significant changes in mouse body weight (**Figure 3C-D**). In addition, the xenograft tumor cells remained sensitive to LGK974 treatment (100 nM) after being harvested and re-cultured *in vitro* (**Figure S3A**), suggesting minimal resistance developed during the *in vivo* tumor progression.

Single cell RNA-seq (scRNA-seq) analysis was further performed to assess the heterogeneity in KPT and KPTN-derived tumor tissues (**Figure S3B-C**). Clustering of 4985 human cells revealed nine shared populations, with clusters 7-9 predominantly present in KPTN tumors (**Figure 3E-F, Supplementary Table 5**). These clusters overexpressed genes involved in ECM remodeling (*MMP7*, *ITGAV*, *COL4A1*, *COL1A1*, *FN1*, *VCAN*, etc.) and YAP signaling (*ANKRD1*) (**Figure 3G**). Notably, they inversely correlated with classical PDAC gene signature while positively associated with basal subtype gene signature (**Figure 3H**), linking *NF2* loss to invasive progression in acinar-derived PDAC. Additionally, projection of bulk RNA-seq KPTN signatures onto scRNA-seq data linked *NF2* loss-induced transcriptomic changes to clusters 7-9. The gene signature of clusters 7-9 is highly expressed in TCGA basal PDAC samples and high-grade samples (**Figure S3D, Supplementary Table 6**). These observations together suggested that *NF2* loss triggers transcriptomic changes in acinar-derived tumors which are associated with a more aggressive phenotype resembling clinical PDAC samples.

Analysis of mouse cancer-associated fibroblasts (CAFs) from tumor tissues revealed a total of 8 clusters of cells (**Figure 3I-J, Supplementary Table 5**). Interestingly, fibroblasts from KPTN tumors were enriched with populations (clusters 6-8) highly expressing *Acta2*, *Spp1* and *Sdc1*, which were associated with a tumor promoting fibroblast phenotype (**Figure 3K**) (25–27). It has been established that TGFβ signaling in the PDAC tumor microenvironment (TME) contributes to the activation of myofibroblast-like CAFs (myCAFs) to foster a more aggressive PDAC phenotype (28). Here we observed higher *TGFβ1* and *Tgfβ1* expression in human tumor cluster 9 and mouse fibroblast cluster 8, respectively, which are both enriched populations in KPTN samples **(Figure 3L**). Consistently, cell-cell communication analysis also inferred an enhanced TGFβ signaling from these 2 clusters (**Figure 3M**), which likely contributes to a tumor-promoting microenvironment in KPTN tumors. In comparison, mouse fibroblasts from KPT tumors were enriched with cells highly expressing tumor-suppressing inflammatory CAFs (iCAF) markers, including *Has1*, *Ccl2*, *Il6* and *Cxcl2* (clusters 1-3) (**Figure 3N, S3E**) (29). Additionally, cells in mouse clusters 1-3 were also found to highly express a recently reported complement-secreting CAFs (csCAF) gene signature (**Figure S3E, Supplementary Table 6**) (30).

### 4. Genetic predisposition primes acinar cells for enhanced cell fitness in the tumor environment

While organoids are widely used in cancer studies (31), their ability to mimic the *in vivo* TME remains unclear. To address this, we performed RNA-seq analysis on KPT and KPTN organoids and revealed 1009 upregulated and 660 downregulated genes in KPTN cells compared to their KPT counterparts (fold change > 2, adjusted P value < 0.05) (**Figure 4A, Supplementary Table 4**). Comparing these with the 683 KPTN tumor-upregulated genes (as described in Figure 3a) revealed 130 consistently upregulated genes (**Figure 4B**), including ECM components (*COL4A1*, *COL4A2*, *COL12A1*, *VCAN, etc.*) and YAP targets (*ANKRD1*, *AXL*, *CYR61*, *F3*) (32). In comparison, other upregulated genes in KPTN samples were found to be context specific. The 553 tumor specifically upregulated genes were enriched in hypoxia, glycolysis, EMT, focal adhesion and proliferation pathways, reflecting a state shift to adapt to the hostile *in vivo* environment. Together, these findings suggested that *NF2* inactivation may trigger intrinsic transcriptomic alterations that prime acinar cells for enhanced fitness in the tumor microenvironment.

**Figure 4.**
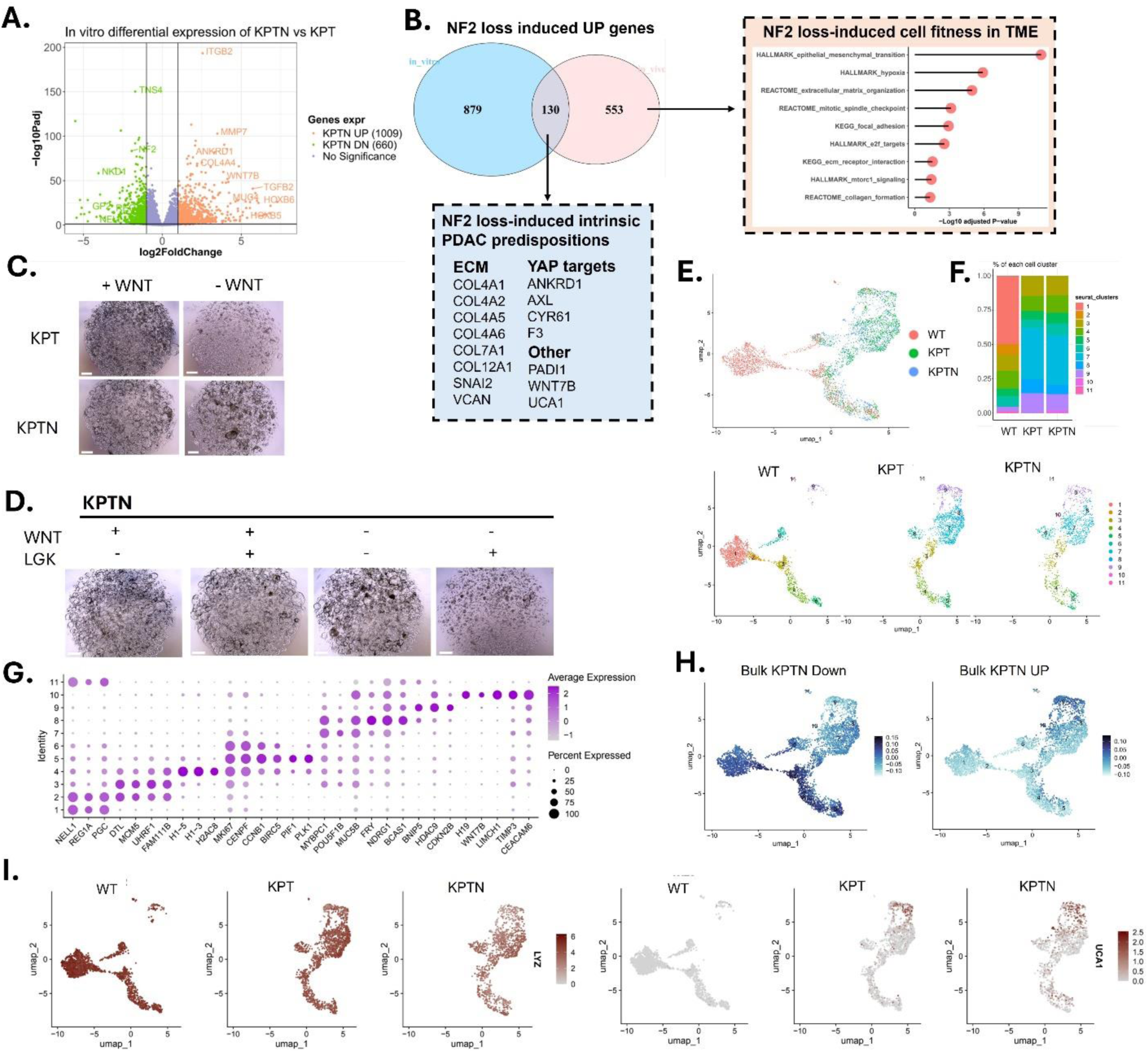
Genetic predisposition primes acinar cells for enhanced cell fitness in the tumor environment. **A.** Volcano plot of differentially expressed genes in KPTN vs KPT *in vitro* organoids (n = 6 for each genotype). **B.** Overlap of the upregulated genes in KPTN *in vitro* cultures and KPTN xenograft tumors (as compared with their KPT analogs, respectively). Boxed blue area: representative genes in the 130 overlapping genes. Boxed red area: overrepresentation analysis of the 553 genes that are only upregulated in KPTN tumors, but not in the *in vitro* culture. **C.** Representative images of KPT and KPTN *in vitro* culture incubated with or without WNT3a supplement. Scale bar = 500 µm. **D.** Representative images of KPTN *in vitro* culture incubated with or without WNT3a supplement or LGK974. Scale bar = 500 µm. **E.** UMAP of all cells from wild type, KPT and KPTN *in vitro* cultures in the single cell RNA seq analysis. **F.** Percentage composition of each cell population as shown in **E**. **G.** Expression of indicated feature genes in each cell population as shown in **E**. **H.** Expression level of indicated gene modules in cell populations shown in **E**. The Bulk KPTN UP and Bulk KPTN DOWN genes were curated from the analysis shown in **A**. **I.** Expression level of indicated genes in cell populations shown in **E**.

Interestingly, *WNT7B* upregulation in both KPTN tumors and organoids suggested that it represented an intrinsic, environmental independent feature. To functionally verify the *WNT7B* upregulation, we cultured organoids in WNT3a-free medium and observed reduced growth from KPT organoids while KPTN ones remained unaffected (**Figure 4C**). Additionally, treatment with 100 nM of LGK974 dramatically suppressed KPTN organoid growth in WNT3a-free medium but not in WNT3a-supplemented medium (**Figure 4D**). This suggested that *NF2* loss drives autonomous WNT7B secretion, resulting in enhanced cell survival in WNT-deprived conditions. Although *GATA6* dysregulation has been linked to elevated WNT ligand expression in PDAC (22), its expression was similar between KPT and KPTN samples (**Figure S4A**), suggesting the existence of alternative mechanisms in regulating WNT expression. These findings establish NF2 inactivation enabling tumor cell survival independent of exogenous WNT signals and promoting tumor progression.

To further probe the heterogeneity in the *in vitro* cultures, we performed scRNA-seq on wild-type (WT), KPT and KPTN organoids and revealed 11 cell clusters (**Figure 4E-F, Supplementary Table 5**). While all 3 genotypes share proliferating cell clusters (3–6) featuring high expression of cell cycle and DNA replication genes, WT organoids have unique cell populations (**Figure 4E-F**) including 2 major *REG1A*+*PGC*+ cluster 1 and 2 cells, representing an acinar to metaplastic cell transitioning stage (**Figure 4G**). In comparison, the unique populations in engineered cells (clusters 7-10) lost all acinar markers and were associated with gene expression found in cancer cells including *MUC5B* and *CEACAM6* (33, 34). Comparison with bulk RNA-seq data (as shown in Figure 4a) demonstrated that cells in clusters 1-6 highly express KPTN Down-regulated genes, while cells in clusters 8-10 highly express KPTN Up-regulated genes (**Figure 4H**). Pseudotime trajectory analysis predicted that the WT metaplastic acinar cells bifurcate toward proliferating or a cancer-like states (**Figure S4B**). The acinar cells gradually lost the classical PDAC subtype gene signature (e.g. *LYZ*, *TFF2* and *AGR2*) upon acquiring driver mutations in the *in vitro* context, whereas high expression of *UCA1*, a basal PDAC subtype marker, was observed in engineered cells (**Figure 4I, S4C**).

### 5. *NF2* inactivation promotes cell survival under nutrient deprivation via enhanced macropinocytosis

To assess the impact of environmental context on acinar transformation, we compared bulk RNA-seq data of KPTN tumors versus organoid cultures. The analysis revealed significant upregulation of genes involved in the cellular response to starvation, and downregulation of genes responsible for glucose and amino acid metabolism in tumor cells (**Figure 5A-B, Supplementary Table 4**). Those transcriptomic changes reflect metabolic rewiring during tumor progression under nutrient-scarce *in vivo* conditions. Comparison of environmentally induced genes (as shown in Figure 5a) with the genetically induced genes (as shown in Figure 3a) identified a total of 378 genes that seem to be primed by genetic predisposition and reinforced by environmental stress (**Figure S5A**). This list included many genes involved in ECM remodeling, YAP signaling, as well as secreted proteins associated with cancer development.

**Figure 5.**
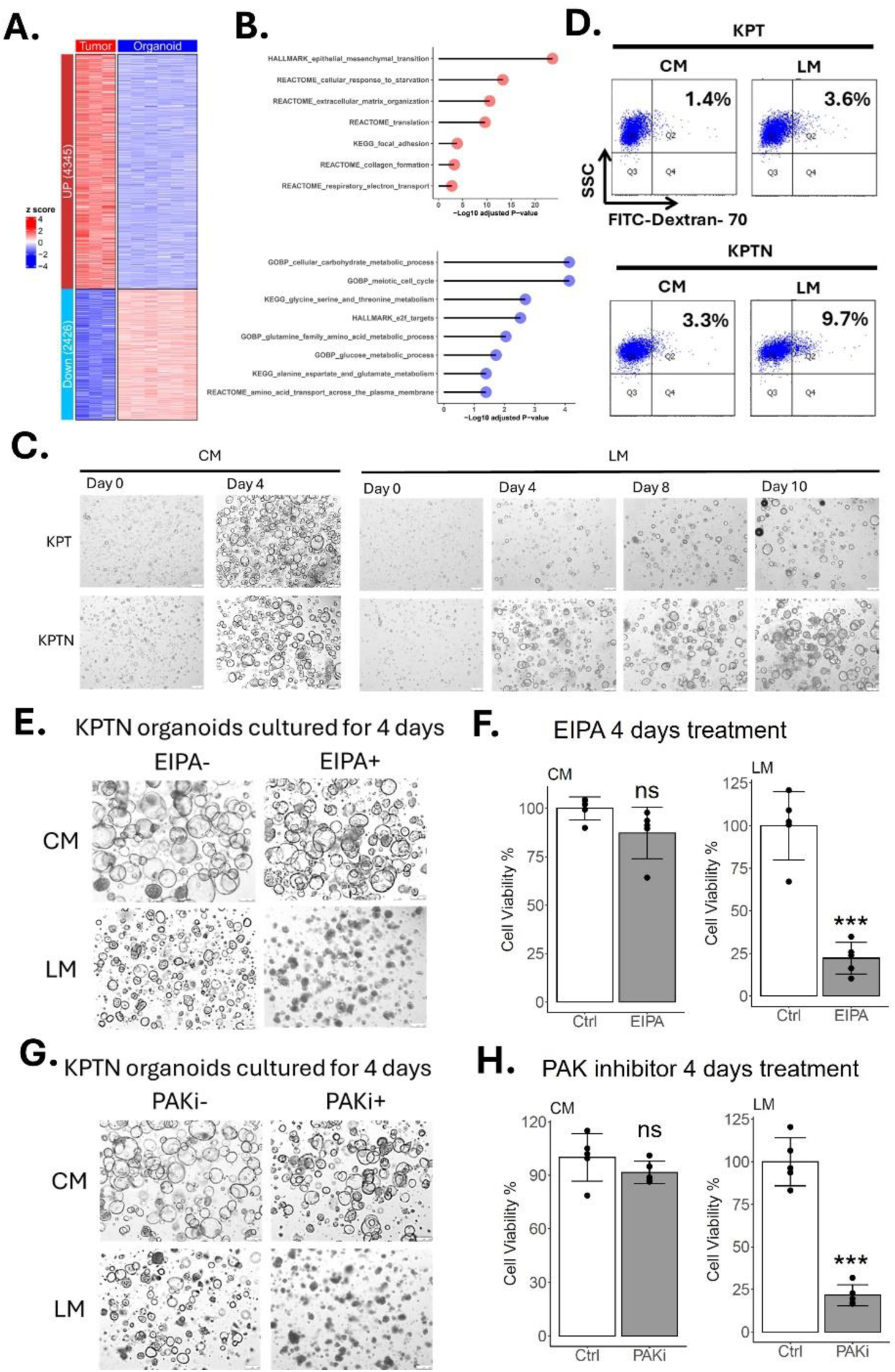
*NF2* inactivation promotes cell survival under nutrient deprivation. **A.** Expression heatmap of differentially expressed genes in KPTN tumor (n = 3) vs KPTN *in vitro* organoid cultures (n = 6). The genes were defined with fold change > 2, adj.p < 0.05. **B.** Overrepresentation analysis for upregulated (in red) and downregulated genes (in blue) in KPTN tumor as shown in **A**. **C.** Representative images of KPT and KPTN organoids cultured in CM or LM for indicated time periods. Scale bar = 250 µm. **D.** Flow cytometry analysis of FITC-Dextran-70 uptake in KPT and KPTN organoids cultured in CM or LM. **E.** Representative images of KPTN organoids cultured in CM or LM and treated with or without EIPA. Scale bar = 250 µm. **F.** Quantification of cell viability of organoids as shown in **E**. Statistical analysis was performed by two-tailed Student’s t test from 3 independent experiments. Error bar represents standard deviation. ns and *** indicate no significance and p-value < 0.001, respectively, compared with vehicle treatment. **G.** Representative images of KPTN organoids cultured in CM or LM and treated with or without PAK inhibitor. Scale bar = 250 µm. **H.** Quantification of cell viability of organoids as shown in **G**.

Analysis of TCGA PDAC data showed that a higher expression score of the induced starvation response genes in KPTN tumors is associated with worse prognosis (**Figure S5B, Supplementary Table 6**). Additionally, it was observed that *NF2* expressions correlate with alterations in multiple cellular metabolism programs in TCGA PDAC patients (**Figure S5C-G**). Based on these findings, we hypothesized that loss of *NF2* contributes to acinar-derived PDAC progression by enhancing cell survival in a nutrient-deprived environment. To test this, KPT and KPTN organoids were cultured in low-nutrient medium (LM) and complete-nutrient medium (CM, see Methods) and monitored for cell growth. While KPT and KPTN cultures exhibited comparable proliferation rates in CM during short-term culture (**Figure 5C**), KPT cells ceased growing in LM after extended culture periods. In contrast, KPTN cells underwent an initial growth arrest but resumed proliferation by day 6-8 in LM.

Macropinocytosis represents one of the crucial mechanisms for cancer cells to combat nutrient starvation (35, 36). It was also reported that Merlin regulates growth factor induced micropinocytosis for membrane receptor recycling (37). Given these facts, we proposed that *NF2* loss could activate macropinosytosis for nutrient scavenging under starvation. To test this, KPT and KPTN organoids were fed with FITC-conjugated dextran-70 for 1 hour, followed by flow cytometry analysis. When cultured in CM, KPTN cells exhibited a higher basal level of dextran uptake compared to KPT cells (3.3% vs. 1.4%) (**Figure 5D**). Importantly, when exposed to LM for 48 hours, dextran uptake was detected in 9.6% of KPTN cells compared to 3.6% of KPT cells, suggesting an enhanced activation of macropinocytosis in KPTN cells under nutrient deprivation. To further examine the contribution of macropinocytosis to cell survival in low-nutrient environments, we cultured the KPTN organoids in CM or LM with or without supplementation of EIPA, a macropinocytosis inhibitor (35, 38). Results showed that EIPA treatment (2.5 µM) dramatically reduced the survival of KPTN cells in LM, while having only minor effect in CM (**Figure 5E-F**). This observation suggested that, in addition to previously reported growth factor-induced macropincytosis, loss of *NF2* also primed cells for macropinocytosis activation upon exposure to nutrient deprivation stress.

Loss of function in *NF2* has been reported to trigger p21-activated kinases (PAK) activation, which was shown to promote macropinocytosis in cancer cells (39, 40). To test whether PAK can mediate macropinocytosis activation upon *NF2* loss, KPTN organoids were cultured in CM or LM with or without FRAX597, a PAK inhibitor (41). Results showed that treatment of FRAX597 at 15 nM completely abolished KPTN survival in LM, while only moderately affecting cell fitness in CM (**Figure 5G-H**). This finding demonstrated that PAK is partially responsible for *NF2* loss-enhanced cell fitness in low-nutrient environments, likely by promoting macropinocytosis.

### 6. Genetic events and environmental cues orchestrate to induce therapeutical resistance

Resistance to therapeutic treatment has been a major obstacle for improving PDAC outcomes. Here we observed that the increased driver mutation burden in engineered acinar cells resulted in enhanced resistance to RMC7977 (at 10 nM to 1 µM), a recently characterized pan-RAS inhibitor (5), highlighting the contribution of genetic predisposition to drug resistance (**Figure S6A**). Interestingly, it was reported that different *in vitro* culture environments could alter the cell states and affect drug responses in patient-derived PDAC organoids (42). We reasoned that the KPTN organoids survived in nutrient-deprived environment may undergo a phenotypic shift, which could result in further altered drug responses. To test this, KPTN cells were allowed to be fully acclimated in LM for 10 days, then treated with gemcitabine and RMC7977 at various doses (**Figure S6B**). The cells continuously cultured in CM were subjected to the same treatment and served as controls. Results showed that while both gemcitabine and RMC7977 were lethal to KPTN organoids at 50 nM, the LM-acclimated KPTN cells exhibited only limited responses (**Figure 6A-B, S6C-D**).

**Figure 6.**
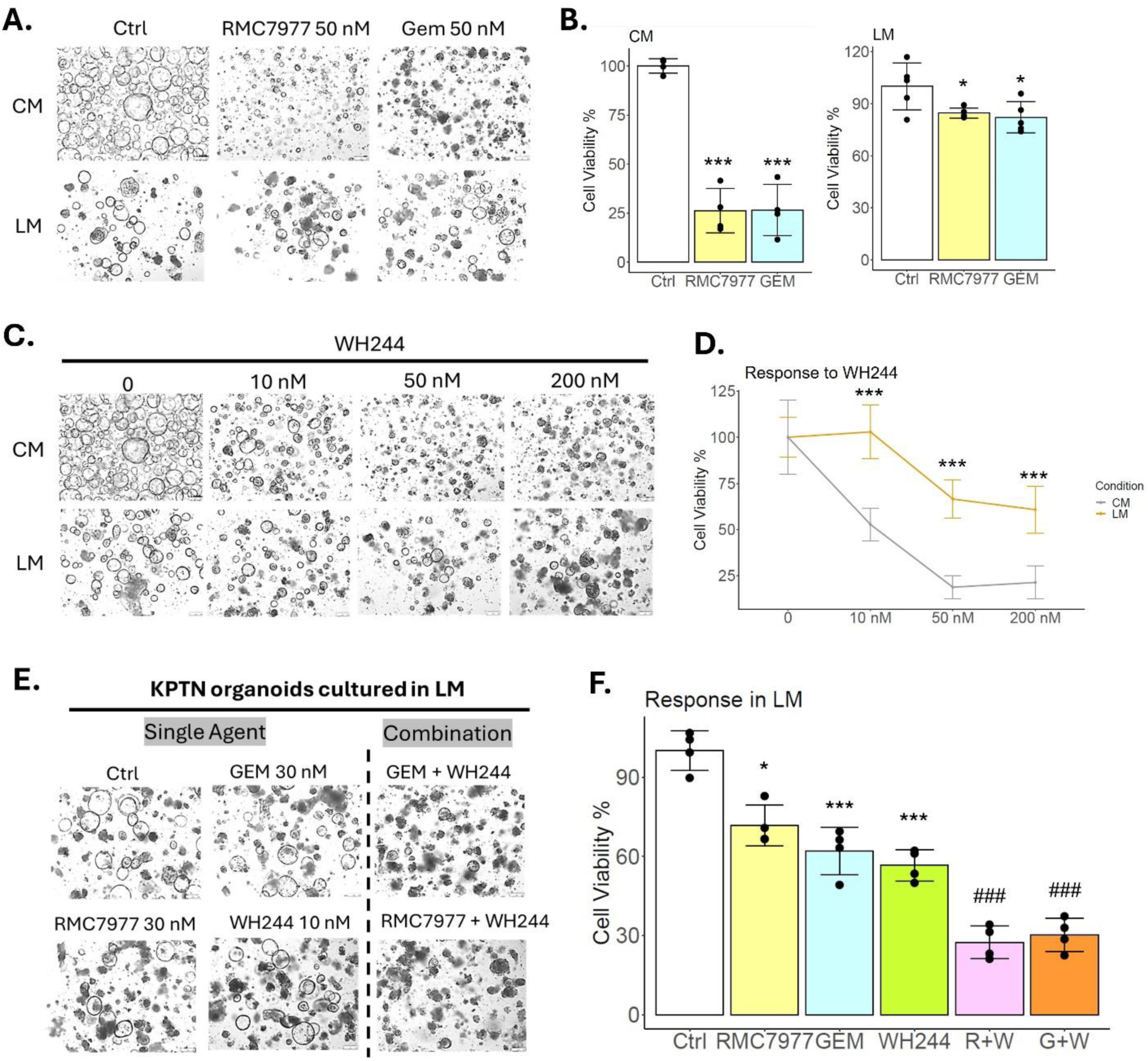
*NF2* loss and nutrient starvation cooperate to induce therapeutic resistance. **A.** Representative images of KPTN organoids cultured in CM or LM and treated with RMC 7977, Gemcitabine or vehicle control. Scale bar = 250 µm. **B.** Quantification of cell viability shown in **A**. Statistical analysis was performed by two-tailed Student’s t test from 3 independent experiments. Error bar represents standard deviation. *, *** indicate p-value < 0.05 and 0.001, respectively, compared with vehicle treatment. **C.** Representative images of KPTN organoids cultured in CM or LM and treated with WH244 or vehicle control. Scale bar = 250 µm. **D.** Quantification of cell viability as shown in **C**. Error bar represents standard deviation. *** indicates p-value < 0.001 between CM versus LM at the same dosage. **E.** Representative images of KPTN organoids cultured in LM and treated with indicated single or combination regime or vehicle control. Scale bar = 250 µm. **F.** Quantification of cell viability of organoids as shown in **E**. Error bar represents standard deviation. *, *** indicate p-value < 0.05 and 0.001, respectively, compared with vehicle treatment. ### indicate p-value < 0.001 compared with single agent treatment.

Dysregulation of apoptosis has been one of the cancer hallmarks and is linked to therapeutic resistance in cancer (43, 44). To test whether inducing apoptosis can reverse the observed drug resistance, we tested a recently developed Bcl-2/Bcl-xL dual degrader (WH244) (45) in KPTN organoids cultured in LM or CM. While WH244 alone showed limited efficacy against LM-acclimated KPTN organoids (**Figure 6C-D**), the combination of WH244 with gemcitabine or RMC7977 led to an improved inhibitory effect compared to single treatment (**Figure 6E-F**). Taken together, our findings revealed that *NF2* inactivation enhances cell survival and therapy resistance under nutrient deprivation likely through metabolic rewiring and macropinocytosis activation.

## DISCUSSION

While *KRAS*, *TP53*, *CDKN2A* and *SMAD4* are the most frequently mutated genes in PDAC, most tumors do not harbor all these mutations, suggesting the existence of other driver mutations. In this study, we employed our unique early human PDAC model combined with CRISPR screening to identify additional tumor suppressor driver mutations that may offer novel treatment options which led to the discovery of *NF2* inactivation as a potent driver of acinar-derived PDAC. Our results demonstrated that *NF2* loss facilitated the aggressive progression of human pancreatic acinar cell-derived PDAC. Mechanistically, *NF2* inactivation induced intrinsic PDAC-associated transcriptomic changes, including alterations in ECM components and elevated niche factor expression, which predispose cells for tumor initiation and progression. Additionally, *NF2* loss primed acinar cells for enhanced cell fitness in hostile environments, such as nutrient deprivation and therapeutic stress.

RNA-seq analysis and validation experiments confirmed that *NF2* inactivation intrinsically upregulated WNT7B expression and secretion, which can be targeted to suppress early PDAC development. While *GATA6* dysregulation has been linked to elevated WNT ligand expression in PDAC patient derived organoids (22), we observed no significant changes in GATA6 expression in KPTN cells, suggesting alternative regulatory mechanisms for WNT upregulation. In addition, *NF2* has been reported to regulate WNT signaling downstream effectors through a FOXM1/β-Catenin axis independent of WNT ligand (46). Our observation of a direct *WNT7B* upregulation upon *NF2* loss provides distinct evidence to support that *NF2* regulates WNT signaling through multiple mechanisms.

There have been concerns as to what extent organoid culture systems can recapitulate the tumor microenvironment in cancer research. It was reported that a state shift in PDAC organoids can be achieved by simply manipulating culture medium components (42), highlighting the flexibility of experimental organoid models. Our data showed that organoid culture can serve as a complementary system to capture intrinsic transcriptomic predisposition driven by genetic alterations. By comparing our *in vitro* and *in vivo* screen outcome as well as RNA-seq data, we demonstrated that, a significant upregulation of genes involved in the cellular response to starvation was observed in tumor cells, while higher expressions of genes responsible for glucose and amino acid metabolism was found in organoids. These findings allowed us to establish a modified low nutrient medium to mimic *in vivo* condition, which revealed an enhanced activation of macropinocytosis by *NF2* inactivation under nutrient stress.

Cancer drug resistance remains a major obstacle to achieving long-term therapeutic success, driven by a complex interplay of genetic, epigenetic, and microenvironmental factors. In this study, we observed that an increased driver mutation burden in engineered acinar cells led to enhanced resistance to a pan-RAS inhibitor, underscoring the role of genetic predisposition in drug resistance. Furthermore, we demonstrated that KPTN organoids adapted to low-nutrient conditions and exhibited resistance to multiple drugs, suggesting that nutrient stress induces a phenotypic shift that exacerbates therapeutic resistance. While many drugs show efficacy against cancer cells and xenograft tumors in preclinical models, they often fail in patients (47), likely due to the lack of a fully recapitulated tumor microenvironment and *in vivo*-like conditions. Our model system addresses this limitation by providing a robust platform to mimic *in vivo* settings. Together, our results emphasize the interplay between genetic alterations and environmental stress in driving therapeutic resistance and suggest that targeting both intrinsic and extrinsic pathways may be necessary to overcome resistance in PDAC.

Previous studies in mice have demonstrated that PDAC derived from different cells of origin may undergo distinct pathological mechanisms (2, 3). While the present study is focused on pancreatic acinar cells, we will take advantage of our unique human PDAC model in future studies to investigate PDAC progression in the ductal lineage. In addition, one limitation of this work is that all the xenograft tumors were generated subcutaneously in immunodeficient mice for efficient screening, which cannot recapitulate the full spectrum of PDAC microenvironment. To address this, we will consider employing humanized mouse models in future studies to enable investigation with the presence of immune components.

In summary, our unique system combining 3D organoid culture of primary human pancreatic cells with CRISPR screen technique serves as an innovative platform to identify additional genes important for PDAC progression. By employing this platform, we identified *NF2* inactivation as a potent driver for human pancreatic acinar transformation, promoting the aggressive progression of acinar-derived PDAC by inducing intrinsic changes to prime transformed cells for an enhanced cell fitness under nutrient deprivation and therapeutic stress. We also identified a list of previously underappreciated tumor suppressor mutations which may provide a selective growth advantage during acinar transformation, warranting further investigation. The discovery of these new drivers in early PDAC development will provide opportunities to explore effective treatment strategies.

## METHODS AND MATERIALS

### Sex as a biological variable

For experiments involving human pancreatic tissues, the sex of organ donors was reported in **Supplementary Table 1** and was not considered as a biological variable. The tissues used in this study were determined based on the availability of resources to provide biological replicates for sufficient statistical power.

For animal experiments, we examined male and female mice, and similar findings are reported for both sexes. The sex was not considered in the study design as no evidence for the impact of sex on the subcutaneous tumorigenesis of pancreatic cancer cells.

### Isolation of human primary pancreatic acinar cells

Human islet-depleted pancreatic exocrine cell fractions were purchased from Prodo Laboratories, Inc (Aliso Viejo, CA). The cells were freshly collected from organ donors deceased due to acute traumatic or anoxic death (**Supplementary Table 1**) and shipped overnight to UT Health San Antonio. The pancreatic acinar cells were isolated by flow sorting as previously described(14). Briefly, exocrine tissue cells were incubated with FITC-conjugated UEA-1 (0.25 μg/ml, Vector Laboratories, Newark, CA, FL-1061-5) for 10 min at 4 °C. After washing with PBS, the cells were digested by incubation with TrypLE^TM^ Express (Life Technologies, Grand Island, NY, 12605-028) for 5-8 min at 37 °C. Cells were collected by centrifugation and washed with FACS buffer (10 mM EGTA, 2% FBS in PBS). Cells were then stained with Pacific blue-conjugated anti-CLA (BioLegend, San Diego, CA, 321308) and anti-7AAD (BioLegend, 420404) for 15 min at 4 °C. Cell pellets were collected by centrifugation and washed with PBS. Flow sorting was performed using a FACSAria^TM^ II (BD Biosciences, San Diego, CA) and the acinar cells were collected in 100% FBS. The collected cells were washed with serum-free Advanced DMEM/F-12 media (Life Technologies, Grand Island, NY, 12634-010) for further process.

### 3D organoid culture and chemical treatment

∼0.5×10^6^ freshly sorted acinar cells were resuspended in 10 µL ice cold Type 2 Cultrex RGF Basement Membrane Extract (BME, R&D Systems, Minneapolis, MN, 3533-010-02P) and were placed in the bottom of a pre-warmed 24 well plate. After solidification at 37 °C for 10 minutes, 500 µL of organoid growth media were added into the well. The media was composed of serum free advanced DMEM/F-12 media (Life Technologies, 12634-010) supplemented with Wnt3a conditioned medium (50%), recombinant human R-Spondin 1 (500 ng/ml, R&D Systems, 4645-RS), Noggin (200 ng/ml, R&D Systems, 6057-NG), FGF10 (100 ng/ml, R&D Systems, 345-FG), EGF (50 ng/ml, R&D Systems, 236-EG), PGE II (1 nM, Fisher Scientific, Waltham, MA, 22-961-0), A83-01 (0.5 µM, R&D Systems, 2939), Nicotinamide (10 mM, Thermo Fisher Scientific, 48-190-7100GM), Penicillin/Streptomycin (1 mM, Sigma-Aldrich, St. Louis, MO, P4333), HEPES (10 mM, Life Technologies, 15630), GlutaMax^TM^ (1X, Life Technologies, 35050).

Modified organoid growth media was prepared as follows and used as indicated in the corresponding results section. The complete nutrient media (CM) was prepared similarly as organoid growth media, except that Wnt3a supplement is absent. The low nutrient medium (LM) was prepared by supplementing basic DMEM (no glucose, no glutamine, Thermo Fisher Scientific, A1443001) with 5% CM. For drug response tests, all the chemicals were prepared as stock solutions followed by dilution in appropriate culture medium to reach final working concentrations. Chemicals used in this study including LGK974 (MedChemExpress, HY-17545), EIPA (MedChemExpress, HY-101840), FRAX597 (MedChemExpress, HY-15542A), RMC-7977 (Chemgood, C-1010), Gemcitabine (ThermoFisher, AC461060010), WH244 (a generous gift from Dr. Daohong Zhou)(45). Organoid images after chemical treatment were captured using an EVOS XL Core or Leica DMI6000 B microscope.

### Genetic engineering human primary pancreatic acinar cells

Introduction of oncogenic KRAS^G12V^ and CRISPR knockout of *p16* and *p53* were performed as described previously(15). Briefly, a plasmid expressing *KRAS^G12V^* cDNA with mCherry was a generous gift from Seung Kim (Stanford University)(48). CRISPR guide RNAs targeting *p16* and *TP53* were cloned into lentiCRISPR v2 vector (Addgene, Watertown, MA, #52961). CRISPR guide RNA targeting *NF2* was cloned into lentiGuide-Hygro-mTagBFP2 vector (Addgene, #99374). Lentiviruses containing these constructs were packaged in 293T cells by co-transfection with packaging plasmids pMD2.G (Addgene, #12259) and psPAX2 (Addgene, #12260). The primary acinar cells were transduced with lentivirus containing each construct with supplement of polybrene at 10 μg/ml. The transduced cells were then subject to antibiotic and/or functional selections. To select *KRAS^G12V^* expressing cells, G418 at 1000 μg/mL was added into organoid culture medium. For the selection of CRISPR-*p16* transduction, cells were treated with puromycin (1 µg/ml). For the selection of *TP53* knockout, 10 µM nutlin-3 was supplemented into the culture medium. To select *NF2* knockout cells, hygromycin at 300 μg/ml was supplemented into the culture medium. The expression of mCherry in the *KRAS^G12V^* cassette was visualized under a Leica DMI6000 B fluorescence microscope. The mutations in *p16* and *TP53* were confirmed by Sanger sequencing. The knockout of *NF2* at protein level was verified by Western Blot.

sgRNA and primer sequences:

*p16* sgRNA: GGCTGGCCACGGCCGCGGCC

*TP53* sgRNA: ACTTCCTGAAAACAACGTTC

*NF2* sgRNA: GCTTGGTACGCAGAGCACCG

*p16* mutation genotyping forward primer: CGGTCCCTCCAGAGGATTTG

*p16* mutation genotyping reverse primer: TGGAGGCTAAGTAGTCCCAG

*TP53* mutation genotyping forward primer: TGCTGGATCCCCACTTTTCC

*TP53* mutation genotyping reverse primer: GGATACGGCCAGGCATTGAA

### CRISPR knockout library preparation

From publicly available data, we compiled a list of 199 recurrently mutated potential tumor suppressor genes reported in clinical PDAC patients(7–13). A CRISPR knockout sgRNA library was designed to target these potential tumor suppressors containing a total of 796 sgRNAs with 4 sgRNAs for each target gene (**Supplementary Table 2**). The pooled sgRNA oligos were purchased from Twist Bioscience (San Francisco, CA). After PCR amplification, pooled sgRNAs were cloned into lentiGuide-Hygro-mTagBFP2 vector (Addgene, #99374) by enzyme digestion followed by ligation using NEBuilder® HiFi DNA Assembly (New England Biolabs, E2621S). Electroporation was then performed using 100 ng of assembly products to transform 50 µL Endura electrocompetent cells at 1800V, 25uF, 200 ohm using a Bio Rad Gene Pulser. After recovered in 2 mL of recovery medium for 1 h, the transformed bacteria cells were plated in a total of 60 of 100 mm agar plates and incubated at 30 C overnight. A total of ∼1.6 × 10^5^ colonies were harvested from all the plates which gave ∼200 coverage for each individual sgRNA. Plasmid DNA was then extracted using GeneJet Midi prep and subject to next generation sequencing to validate the distribution of each individual sgRNA in the library. The plasmid DNA was used to transfect 293T cells using Lipofectamine 3000 reagent, together with packaging plasmids pMD2.G and psPAX2, for lentivirus production.

### CRISPR knockout screen and data analysis

The pancreatic acinar organoids were transduced with lentivirus containing the CRISPR sgRNA library at a low MOI of ∼ 0.3 followed by antibiotic selection with hygromycin at 300 μg/ml. A successful transduction was verified by confirming the cellular expression of BFP tag from the vector construct using a Leica DMI6000 B fluorescence microscope. The transduced cells were expanded for one passage, a fraction of cells was collected to serve as a reference control. For the *in vivo* screen, the expanded cells were subcutaneously transplanted into the hind flank of NSG mice. The xenograft tumor tissues were collected after 8 weeks followed by DNA extraction using a Genomic DNA Clean & Concentrator-25 kit (Zymo Research, Irvine, CA, D4065). For the *in vitro* screen, the expanded cells were continuously passaged for 8 weeks (7-9 passages), followed by extraction of cellular DNA. The collected DNA containing sgRNA library sequences was subject to a two-step PCR protocol for library amplification and sample indexing, followed by NGS analysis at the Greehey Children’s Cancer Research Institute (GCCRI) Genome Sequencing Facility at UT Health San Antonio. The screen was performed using 4 independently established acinar cultures, with 2 replicates of *in vivo* tumor samples and 1 replicate of *in vitro* culture sample for each. Count summary and statistical analysis of sgRNA enrichment from fastq data was performed using MAGeCK software v0.5.9.5(49). A significant positive enrichment at target gene level was considered with a fold change > 1 and p-value < 0.05 compared with corresponding reference control sample.

### 3D organoid cell viability assay

Cell viability of 3D organoids culture after chemical treatment was assessed using CellTiter-Glo® 3D Cell Viability Assay Kit (Promega, Madison, WI) according to manufacturer’s instruction. Briefly, 8000 cells were resuspended in 5 μl of ice cold BME and placed in the center of each well in a 96 well plate. After solidification at 37 °C for 10 minutes, 100 µL of organoid growth media were added into each well. After 3 days of incubation, the medium was refreshed with supplementation of different chemicals as indicated in the corresponding result section. At the end point, 100 µL of CellTiter-Glo® 3D Cell Viability reagent was added into each well followed by vigorous mixing to induce cell lysis. The plate was sat at room temperature for 25 min and subject to luminescence detection using a BioTek Synergy H1 microplate reader.

### Flow cytometry analysis for BFP and FITC-dextran

KPT and KPTN cells were mixed at 1:1 ratio and embedded in BME for 3D organoid culture. A fraction of the mixed cells was collected at passage 1 to passage 7 for flow cytometry analysis of the BFP signal inserted in the *NF2* sgRNA construct using a BD^®^ LSR II flow cytometer. KPT and KPTN monoculture were subject to the same analysis as controls.

For dextran uptake analysis, KPT and KPTN cells were cultured in complete nutrient medium for 4 days to allow organoid formation. On day 4, the organoids were exposed to low-nutrient medium for 1 day to induce starvation responses. A control group was maintained in fresh complete nutrient medium. Half of the medium (∼200 µL) from each well was collected, and FITC-conjugated dextran 70 (Thermo Fisher Scientific, D1822) was added to reach a final concentration of 1 mg/ml. Dispase was added to the remaining medium (2U/ml) and incubated at 37°C for 20 minutes to release organoids from BME. The organoids were then collected by centrifugation at 200g for 3 minutes. Half of the organoids were kept as a baseline control, while the other half was resuspended in dextran 70-containing medium and incubated at 37°C for 30 minutes. The organoids were then washed twice with PBS and dissociated into single cells for flow cytometry analysis using a BD^®^ LSR II flow cytometer.

### Animal experiment

All animal experiment protocols were reviewed and approved by the Institutional Animal Care and Use Committees at UT Health San Antonio (20130023AR) and performed in accordance with relevant guidelines and regulations. The maximal xenograft tumor size in each mouse did not exceed 2 cm at the largest diameter as permitted by institutional guidelines. For all the animal experiments, six-eight week old female and male NSG (NOD.Cg-Prkdc scid Il2rg tm1Wjl/SzJ) mice were employed which were purchased from The Jackson Laboratory (Strain #:005557). The mice were housed at a maximum of five per cage in a pathogen-free system with water and food ad libitum, with 12 hours day/night cycle at 20-25 °C and 50-60% humidity. Detailed study design and sample size of animal studies were described below, which were determined based on the availability of resources to provide at least 4 biological replicates for a sufficient statistical power. No criteria used for including and excluding animals during the experiment. No exclusions of any animals or data points in the data analysis. Randomisation was not done in this study, and confounders were not controlled. When comparing samples from different groups, we used paired samples from the same culture origin, and made sure to include multiple biological replicates to minimize potential heterogeneity. The authors who were responsible for animal experiments as well as following data analysis were aware of the group allocation at all stages of the experiment.

For *in vivo* CRISPR screen experiments, detailed study design was described previously under “CRISPR knockout screen and data analysis” sub-section. Briefly, a total of 8 NSG mice were subject to xenograft transplantation (4 independently established acinar cultures, with 2 replicates each). Tumor xenografts were established by subcutaneously injecting ∼1×10^6^ engineered cells (suspended in 100 µL of 50% BME/50% Advanced DMEM/F-12) into the hind flank of mice. The xenograft tumor tissues were collected after 8 weeks followed by DNA extraction followed by NGS analysis. The enrichment of any sgRNA in tumor tissue DNA was identified by comparing with paired control as described previously. For functional verification of the effect from *NF2* knockout on tumor development, a total of 45 mice were subject to subcutaneous transplantation of ∼1×10^6^ engineered cells (n = 4-7 for each of the 4 independent cultures and each of the 2 genotypes, as indicated in Figure 2c). After 8 weeks, the mice were euthanized to collect the tumor tissues for further histological analysis. The difference in tumor size was compared between KPTN tumors versus KPT counterparts from the same culture origin. Tumor volume was calculated as: V = L×W^2^/2. To assess the effect of LGK974 on KPTN tumor growth, a total of 8 mice received transplantation of ∼1×10^6^ KPTN cells at both sides of hind flank. After 1 month to allow tumor development, the tumor bearing mice were separated into 2 groups (4 mice for each group) and received a daily dose of LGK974 (10 mg/kg, oral gavage) or vehicle control for 40 days. Tumor size and mice body weight were recorded every 5 days. At the endpoint, the mice were euthanized to collect tumor tissues for further analysis.

### Western blot

Protein expression in the organoid culture was detected by Western Blot following a standard protocol. Briefly, the cells were harvested to collect cell lysates. ∼20 ug protein from cell lysates was loaded into each lane of a 10% SDS-PAGE gel followed by electrophoresis at a voltage of 90 V. The proteins on the gel were then transferred to a PVDF membrane at constant voltage of 80 V for 15 h on ice. After transferring, the membranes were incubated in blocking solution (3% BSA in TBST) with continuous rock for 60 min, then incubated with primary antibody overnight at 4°C. The membranes were then washed and incubated with HRP-conjugated secondary antibody for 2 h at room temperature. The membranes were subsequently washed and incubated with ECL western blot substrates for 2 min, followed by signal capture using an Amersham Imager 600. All primary and secondary antibodies used are listed below:

rabbit anti-human NF2 (Abclonal, #A0739), dilution 1:500;

mouse anti-human GAPDH (Santa Cruz, #sc-32233), dilution 1:500;

mouse anti-rabbit IgG-HRP (Santa Cruz, #sc-2357), dilution 1:1000;

goat anti-mouse IgG-HRP (Santa Cruz, #sc-2005), dilution 1:1000.

### H&E and IF staining

Freshly collected xenograft tumor tissues were fixed to prepare paraffin-embedded block. Tissue sectioning, H&E and alcian blue staining were performed at the histology laboratory at UT Health San Antonio following standard protocols. For IF staining, paraffin-embedded tissue section were deparaffinized, rehydrated, and submerged in 200 °C heated R-Universal Epitope Recovery Buffer solution (Electron Microscopy Sciences, Hatfield, PA, #AP0530) for 30 minutes.

After sitting at room temperature for 20 minutes, sections were permeabilized using 0.5% PBST (0.5% Triton X-100 in PBS) for 10 minutes and blocked with 5% donkey serum in 0.1% PBST for 60 minutes at room temperature. Sections were then incubated with primary antibodies diluted in blocking solution at 4 °C overnight. Sections were subsequently incubated with fluorescent-tagged Alexa Fluor secondary antibodies diluted in 5% blocking solution for 1 hour at room temperature. Additionally, sections were incubated with DAPI (1:1000, Invitrogen, Carlsbad, CA, P36935) for 4 minutes at room temperature. Finally, sections were covered with a drop of VectaShield Vibrance Antifade Mounting Medium (Vector Laboratories, Inc., Burlingame, CA, H-1700). All histology and fluorescence images were captured using a Leica DMI6000 B microscope and a compatible software (Leica Microsystems, Buffalo Grove, IL). All primary and secondary antibodies used are listed below:

Mouse anti human STEM121 (Takara Bio, #Y40410), dilution 1:200;

Mouse anti human/mouse/rat KRT19 (DSHB, #Troma-III), dilution 1:50;

Mouse anti human/mouse/rat YAP (63.7) (Santa Cruz, # sc-101199), dilution 1:50;

Alexa Fluor® 488-conjugated AffiniPure Donkey Anti-Mouse IgG (H+L) (Jackson ImmunoResearch, #715-545-150), dilution 1:250.

### Bulk RNA seq library preparation and data analysis

Cell pellets of cultured organoids were collected and stored in Trizol for total RNA extraction using Zymo Direct-zol RNA Miniprep Kit (Zymo Research, Irvine, CA, R2063) following manufacturer’s instructions. Tumor tissues were flash frozen in liquid nitrogen and ground into fine powder with the addition of Trizol, followed by RNA extraction using Zymo Direct-zol RNA Miniprep Kit. Indexed cDNA libraries were prepared using the Illumina® Stranded mRNA Prep Ligation kit (Illumina, San Diego, CA, 20040532) following manufacturer’s instructions. The cDNA libraries were then submitted to the GCCRI Genome Sequencing Facility at UT Health San Antonio for high throughput sequencing analysis using Illumina HiSeq 3000 or NovaSeq 6000 System.

Raw sequencing data were aligned to reference genome GRCh38 or GRCm38 using TopHat 2.1.1. Gene expression reads were quantified using HTSeq 0.11.1. For tumor tissue samples, in order to separate human and mice transcripts, the raw reads were aligned to a concatenated human and mouse genome. Reads for which multiple alignments shared the top score were discarded to remove those aligned to both the human and mouse genome(50). Differential expression analysis of the read counts from aligned RNA seq data was performed using DESeq2 1.36.0 package in R software. A pre-filtering process was applied for each analysis to remove genes with low read count, i.e. only retained genes with > 10 reads in at least half of the samples of at least one comparing group. The differentially expressed genes (DEGs) were defined as fold change > 2, and Benjamin-Hochberg (BH) adjusted p-value < 0.05. Gene expression heatmaps were generated using z-score of transformed/normalized read count as indicated in each figure. Overrepresentation analysis for lists of genes were performed using the clusterProfiler 4.4.4 package in R with default settings. A significant enrichment was considered with multiple-test adjusted p-value < 0.05.

### Single cell RNA seq library preparation and data analysis

Single cell RNA seq samples were prepared using an Evercode^TM^ Fixation V2 kit and Evercode^TM^ WT V2 Kit (Parse Biosciences, Seattle, WA) following manufacturer’s instructions. Briefly, organoid culture or tumor tissues were dissociated into single cells by enzyme digestion and flow sorting. The single cells were subject to a fixation procedure using the Evercode^TM^ Fixation V2 kit followed by a split-pool combinatorial barcoding protocol in 96 well format using the Evercode^TM^ WT V2 Kit. The barcoded cDNAs were equally dispersed into 8 indexed sub-libraries and were submitted for high throughput sequencing analysis using Illumina NovaSeq X plus System.

Processing of the raw sequencing reads including alignment, demultiplexing and separation of human and mice reads in the scRNA seq data were performed using a ParseBiosciences-Pipeline software v1.1.2 which was designed to be compatible with the sample preparation protocol. Reference genomes GRCh38 and GRCm38 were used for alignment. The subsequent analysis was performed using Seurat v.5.0.1 in R software. A quality control step was performed to remove low quality cells, including possible cell debris (nFeature < 200), cell doublets (nFeature > 9,000 for organoid samples and nFeature > 5,000 for tumor samples), and cells with excessive cellular stress (> 15% human mitochondrial gene or > 10% mouse mitochondrial gene expression). The filtered Seurat object was subjected to a series of standard Seurat commands to identify different cell populations present in the data set. For *in vitro* organoid samples, an extra “harmony integration” step was performed before downstream analysis to remove batch effects likely introduced during sample preparation. For *in vivo* tumor samples, the combined Seurat object was subset to contain human or mouse cells only for downstream analysis. Expression scores of a list of gene signature were calculated using “AddModuleScore” command in Seurat. Single cell trajectory analysis was performed using Monocle3 v.1.3.4 in R software with default setting. The Seurat object containing all organoid cells were passed into Monocle3 to infer their cell-type transition states, by setting cluster 1 as the root. Cell-cell communication analysis was performed using CellChat v2.1.2 in R software with default setting.

### Kaplan-Meier plot of TCGA data

Clinical prognosis of gene expression was assessed using TCGA PDAC patient data. For prognosis analysis of a single gene, the patients were grouped by median gene expression using RNA seq data followed by Kaplan-Meier plot with survival data. For prognosis of a group of genes, the gene signature expression score of each patient was calculated using GSVA v1.52.3 from their RNA seq data(51). Then the patients were grouped by median expression score followed by Kaplan-Meier plot. The KM plot was generated using survival v3.7 in R environment.

### Statistical analysis

Statistical analyses of CRISPR screen enrichment were performed using MAGeCK algorithm. All the bioinformatic analyses of high throughput RNA sequencing data were performed using the R packages described in corresponding method sections and figure legends. Quantitative analysis of experimental results between 2 groups was performed using two tailed Student’s t test. The sample size in each experiment was described in corresponding result sections and figure legends.

### Study approval

All experiments in this study using primary human pancreatic tissues were reviewed by the UT Health San Antonio Institutional Review Board (IRB). The patient donors were de-identified, with available information on gender, race, age, weight, height and cause of death. The IRB committee agreed that this study does not require IRB approval because it is either: not human research as defined by DHHS regulations at 45 CFR 46 and FDA regulations at 21 CFR 56; the project does not include non-routine intervention or interaction with a living individual for the primary purpose of obtaining data regarding the effect of the intervention or interaction, nor do the researchers obtain private, identifiable information about living individuals.

All animal experiment protocols were approved by the UT Health San Antonio Institutional Animal Care and Use Committees (protocol #20130023AR) and were performed in accordance with relevant guidelines and regulations.

## Data Availability

A **Supporting Data Values** file was provided which includes all data in the manuscript and supplement represented in graphs and as mean ± standard deviation. The raw and processed high throughput sequencing data generated in this study have been deposited to and will be publicly available at Gene Expression Omnibus under accession numbers **GSE292511**, **GSE292512** and **GSE292513**. TCGA pancreatic cancer patient data are available at GDC data portal [https://portal.gdc.cancer.gov/projects/TCGA-PAAD]. Human reference genome GRCh38 and mouse reference genome GRCm38 are available at [https://genome.ucsc.edu/index.html].

## Author Contributions

Conceptualization: Wang P, Liu J

Data curation: Xu Y, Nipper M, He C, Liu J, Wang P

Formal analysis: Xu Y, Nipper M, Liu J, Dominguez A, He C, Sharkey FE, Khan S, Zhou D, Xu H, Luan Y, Wang P

Funding acquisition: Wang P

Methodology: Xu Y, Nipper M, Liu J

Project administration: Wang P and Liu J

Supervision: Wang P

Writing – original draft: Xu Y, Liu J, Wang P

Writing – review & editing: Xu Y, Nipper M, Zhou D, Zheng L, Luan Y, Liu J, Wang P

## Acknowledgements

This work is supported by the Cancer Prevention and Research Institute of Texas (R1219, P. Wang), NIH/NCI (R21 CA218968, P. Wang, R01 CA237159, P. Wang), NIDDK (R01 DK110361, P. Wang), and the William and Ella Owens Medical Research Foundation (P. Wang). P. Wang is a Cancer Prevention and Research Institute of Texas (CPRIT) scholar. A. Dominguez is supported by T32CA148724 and T32GM148752. The funders had no role in study design, data collection and analysis, decision to publish, or preparation of the manuscript.

The authors acknowledge Dr. Seung Kim (Stanford University) for kindly providing KRAS construct. The authors acknowledge Dr. Junichi Takagi for kindly providing engineered cell line that produced biologically active Wnt/afamin. Part of data was generated in the Flow Cytometry Shared Resource at UT Health San Antonio which is supported by the NCI grant from Mays Cancer Center (P30CA054174), the CPRIT grant (RP210126) and NIH grant (1S10OD030432-01A1). Part of the histology studies were assisted by the Histology and Immunohistochemistry Laboratory at UT Health San Antonio. High throughput sequencing data described in this study was generated in the Genome Sequencing Facility/Mays Cancer Center Next-generation Sequencing Facility, which is supported by UT Health San Antonio, NIH-NCI P30 CA054174 (Cancer Center at UT Health San Antonio) and NIH Shared Instrument grant S10OD030311 (S10 grant to NovaSeq 6000 System), and CPRIT Core Facility Award (RP220662).

## Abbreviation

CAF: cancer-associated fibroblast
CM: complete nutrient medium
DEG: differentially expressed genes
ECM: extracellular matrix
IF: immunofluorescence
KPT: *KRAS*, *CDKN2A*, *p53* triple mutation
KPTN: *KRAS*, *CDKN2A*, *p53, NF2* quadruple mutation
LM: low nutrient medium
PDAC: pancreatic ductal adenocarcinoma
scRNA-seq: single cell RNA-sequencing
sgRNA: single guide RNA
TGFB: transforming growth factor beta

## REFERENCES

1. Siegel RL, Giaquinto AN, Jemal A. Cancer statistics, 2024. 10.3322/caac.21820.

2. Flowers BM, et al. Cell of Origin Influences Pancreatic Cancer Subtype. Cancer Discov. 2021;11(3):660–677.

3. von Figura G, et al. The chromatin regulator Brg1 suppresses formation of intraductal papillary mucinous neoplasm and pancreatic ductal adenocarcinoma. Nat Cell Biol. 2014;16(3):255– 267.

4. Alonso-Curbelo D, et al. A gene–environment-induced epigenetic program initiates tumorigenesis. Nature. 2021;590(7847):642–648.

5. Holderfield M, et al. Concurrent inhibition of oncogenic and wild-type RAS-GTP for cancer therapy. Nature. 2024;629(8013):919–926.

6. Wasko UN, et al. Tumour-selective activity of RAS-GTP inhibition in pancreatic cancer. Nature. 2024;629(8013):927–936.

7. Jones S, et al. Core signaling pathways in human pancreatic cancers revealed by global genomic analyses. Science. 2008;321(5897):1801–1806.

8. Bailey P, et al. Genomic analyses identify molecular subtypes of pancreatic cancer. Nature. 2016;531(7592):47–52.

9. Waddell N, et al. Whole genomes redefine the mutational landscape of pancreatic cancer. Nature. 2015;518(7540):495–501.

10. Biankin AV, et al. Pancreatic cancer genomes reveal aberrations in axon guidance pathway genes. Nature. 2012;491(7424):399–405.

11. Witkiewicz AK, et al. Whole-exome sequencing of pancreatic cancer defines genetic diversity and therapeutic targets. Nat Commun. 2015;6:6744.

12. Roberts NJ, et al. Whole Genome Sequencing Defines the Genetic Heterogeneity of Familial Pancreatic Cancer. Cancer Discov. 2016;6(2):166–175.

13. Cancer Genome Atlas Research Network. Electronic address, Cancer Genome Atlas Research Network. Integrated Genomic Characterization of Pancreatic Ductal Adenocarcinoma. Cancer Cell. 2017;32(2):185–203.e13.

14. Liu J, et al. TGF-β1 promotes acinar to ductal metaplasia of human pancreatic acinar cells. Sci Rep. 2016;6:30904.

15. Xu Y, et al. Reconstitution of human PDAC using primary cells reveals oncogenic transcriptomic features at tumor onset. Nat Commun. 2024;15(1):818.

16. Akanuma N, et al. Paracrine Secretion of Transforming Growth Factor β by Ductal Cells Promotes Acinar-to-Ductal Metaplasia in Cultured Human Exocrine Pancreas Tissues. Pancreas. 2017;46(9):1202–1207.

17. Nipper M, et al. TGFβ and Hippo Signaling Pathways Coordinate to Promote Acinar to Ductal Metaplasia in Human Pancreas. Cells. 2024;13(2):186.

18. Wang W, et al. ARID1A, a SWI/SNF subunit, is critical to acinar cell homeostasis and regeneration and is a barrier to transformation and epithelial-mesenchymal transition in the pancreas. Gut. [published online ahead of print: September 18, 2018]. 10.1136/gutjnl-2017-315541.

19. Thurneysen C, et al. Functional inactivation of NF2/merlin in human mesothelioma. Lung Cancer. 2009;64(2):140–147.

20. Bachir S, et al. Neurofibromatosis Type 2 (NF2) and the Implications for Vestibular Schwannoma and Meningioma Pathogenesis. Int J Mol Sci. 2021;22(2):690.

21. Moffitt RA, et al. Virtual microdissection identifies distinct tumor- and stroma-specific subtypes of pancreatic ductal adenocarcinoma. Nat Genet. 2015;47(10):1168–1178.

22. Seino T, et al. Human Pancreatic Tumor Organoids Reveal Loss of Stem Cell Niche Factor Dependence during Disease Progression. Cell Stem Cell. 2018;22(3):454–467.e6.

23. Tammela T, et al. A Wnt-producing niche drives proliferative potential and progression in lung adenocarcinoma. Nature. 2017;545(7654):355–359.

24. Liu J, et al. Targeting Wnt-driven cancer through the inhibition of Porcupine by LGK974. Proc Natl Acad Sci. 2013;110(50):20224–20229.

25. Öhlund D, et al. Distinct populations of inflammatory fibroblasts and myofibroblasts in pancreatic cancer. J Exp Med. 2017;214(3):579–596.

26. Eun JW, et al. Cancer-associated fibroblast-derived secreted phosphoprotein 1 contributes to resistance of hepatocellular carcinoma to sorafenib and lenvatinib. Cancer Commun Lond Engl. 2023;43(4):455–479.

27. Maeda T, Alexander CM, Friedl A. Induction of syndecan-1 expression in stromal fibroblasts promotes proliferation of human breast cancer cells. Cancer Res. 2004;64(2):612–621.

28. Elyada E, et al. Cross-Species Single-Cell Analysis of Pancreatic Ductal Adenocarcinoma Reveals Antigen-Presenting Cancer-Associated Fibroblasts. Cancer Discov. 2019;9(8):1102– 1123.

29. Oh K, et al. Coordinated single-cell tumor microenvironment dynamics reinforce pancreatic cancer subtype. Nat Commun. 2023;14(1):5226.

30. Chen K, et al. Single-cell RNA-seq reveals dynamic change in tumor microenvironment during pancreatic ductal adenocarcinoma malignant progression. EBioMedicine. 2021;66:103315.

31. Verstegen MMA, et al. Clinical applications of human organoids. Nat Med. 2025;31(2):409–421.

32. Wang Y, et al. Comprehensive Molecular Characterization of the Hippo Signaling Pathway in Cancer. Cell Rep. 2018;25(5):1304–1317.e5.

33. Muilenburg KM, et al. Mucins as contrast agent targets for fluorescence-guided surgery of pancreatic cancer. Cancer Lett. 2023;561:216150.

34. Nakazawa Y, et al. Delivery of a BET protein degrader via a CEACAM6-targeted antibody-drug conjugate inhibits tumour growth in pancreatic cancer models. Nat Commun. 2024;15(1):2192.

35. Kim SM, et al. PTEN Deficiency and AMPK Activation Promote Nutrient Scavenging and Anabolism in Prostate Cancer Cells. Cancer Discov. 2018;8(7):866–883.

36. Wang T, et al. Amino Acid-Starved Cancer Cells Utilize Macropinocytosis and Ubiquitin-Proteasome System for Nutrient Acquisition. Adv Sci Weinh Baden-Wurtt Ger. 2024;11(1):e2304791.

37. Chiasson-MacKenzie C, et al. Merlin/ERM proteins regulate growth factor-induced macropinocytosis and receptor recycling by organizing the plasma membrane:cytoskeleton interface. Genes Dev. 2018;32(17–18):1201–1214.

38. Jayashankar V, Edinger AL. Macropinocytosis confers resistance to therapies targeting cancer anabolism. Nat Commun. 2020;11(1):1121.

39. Kissil JL, et al. Merlin, the product of the Nf2 tumor suppressor gene, is an inhibitor of the p21-activated kinase, Pak1. Mol Cell. 2003;12(4):841–849.

40. Lee S-W, et al. EGFR-Pak Signaling Selectively Regulates Glutamine Deprivation-Induced Macropinocytosis. Dev Cell. 2019;50(3):381–392.e5.

41. Licciulli S, et al. FRAX597, a small molecule inhibitor of the p21-activated kinases, inhibits tumorigenesis of neurofibromatosis type 2 (NF2)-associated Schwannomas. J Biol Chem. 2013;288(40):29105–29114.

42. Raghavan S, et al. Microenvironment drives cell state, plasticity, and drug response in pancreatic cancer. Cell. 2021;184(25):6119–6137.e26.

43. Kim M, et al. SLC5A3 depletion promotes apoptosis by inducing mitochondrial dysfunction and mitophagy in gemcitabine-resistant pancreatic cancer cells. Cell Death Dis. 2025;16(1):1– 11.

44. Low HB, et al. DUSP16 promotes cancer chemoresistance through regulation of mitochondria-mediated cell death. Nat Commun. 2021;12(1):2284.

45. Nayak D, et al. Development and crystal structures of a potent second-generation dual degrader of BCL-2 and BCL-xL. Nat Commun. 2024;15(1):2743.

46. Quan M, et al. Merlin/NF2 Suppresses Pancreatic Tumor Growth and Metastasis by Attenuating the FOXM1-Mediated Wnt/β-Catenin Signaling. Cancer Res. 2015;75(22):4778– 4789.

47. Hay M, et al. Clinical development success rates for investigational drugs. Nat Biotechnol. 2014;32(1):40–51.

48. Lee J, et al. Reconstituting development of pancreatic intraepithelial neoplasia from primary human pancreas duct cells. Nat Commun. 2017;8:14686.

49. Li W, et al. MAGeCK enables robust identification of essential genes from genome-scale CRISPR/Cas9 knockout screens. Genome Biol. 2014;15(12):554.

50. Callari M, et al. Computational approach to discriminate human and mouse sequences in patient-derived tumour xenografts. BMC Genomics. 2018;19(1):19.

51. Hänzelmann S, Castelo R, Guinney J. GSVA: gene set variation analysis for microarray and RNA-seq data. BMC Bioinformatics. 2013;14:7.

