## Supplementary Figures for "*NF2* loss malignantly transforms human pancreatic acinar cells and enhances cell fitness under environmental stress"

Yi Xu *et al.*

This PDF file includes:

- Figure. S1 to S6 and corresponding legends

- Description of Supplementary Tables 1-6

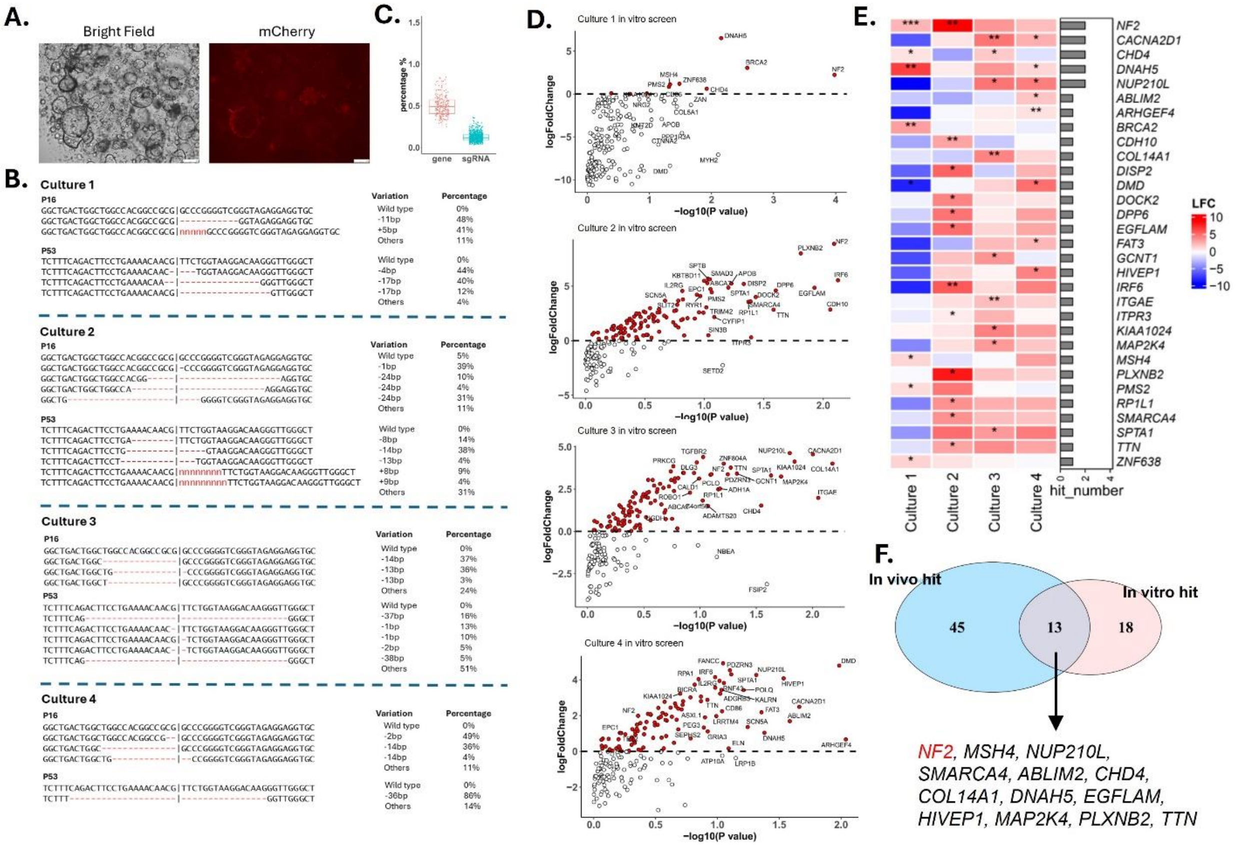


**Figure S1. A.** Bright field and fluorescence images of acinar organoids transduced with lentivirus expressing mCherry tagged KRAS$^{G12V}$ construct, scale bar = 250 µm. **B.** Sanger sequencing analysis of p16 and p53 at the sgRNA target site in 4 independent cultures. **C.** Unbiased distribution of all 796 sgRNAs and corresponding target genes in the library plasmid DNA. **D.** Scatter plot of the enrichment of target genes from *in vitro* screen in 4 independent cultures. The Y axis represents log2 fold change of the sgRNAs distribution of an indicated target gene compared with the control. X axis represents -log10 P value. Positively enriched targets (log2 fold change > 0) in individual replicate were highlighted in red. **E.** Heatmap of all target genes whose sgRNAs were found to be positively enriched (fold change > 1, P value < 0.05) in any of the 4 independent cultures. The color scale represents log2 fold change (LFC). The side bar plot indicates the number of replicates in which each target was found to be enriched. *, **, *** indicate p-value < 0.05, 0.01, 0.001, respectively. **F.** Comparison of the enriched target genes from *in vivo* and *in vitro* screen.

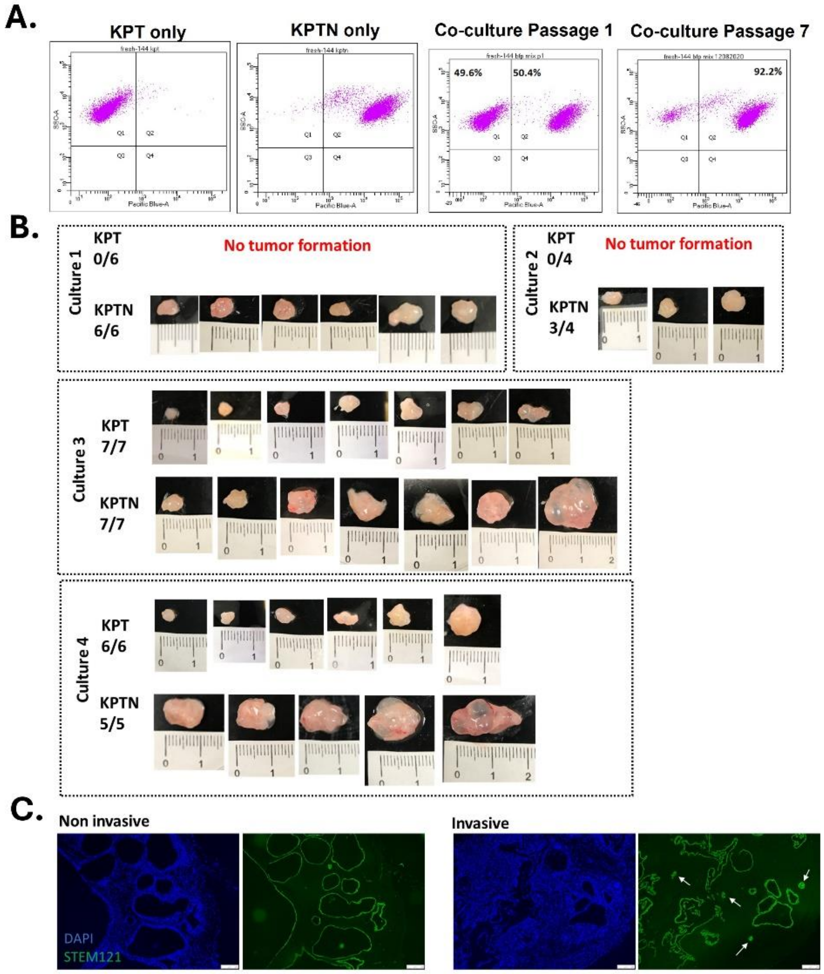


**Figure S2. A.** Flow cytometry analysis of BFP signal in KPT single culture, KPTN single culture or their coculture at 1:1 ratio. **B.** Photos of all xenograft tumors harvested from NSG mice transplanted with KPT or KPTN organoids. **C.** Representative immunofluorescence staining of human specific STEM121 in noninvasive and invasive tumor sections. Cell nuclei was counter stained with DAPI. Scale bar = 50 µm. Arrows indicate invasive lesions.

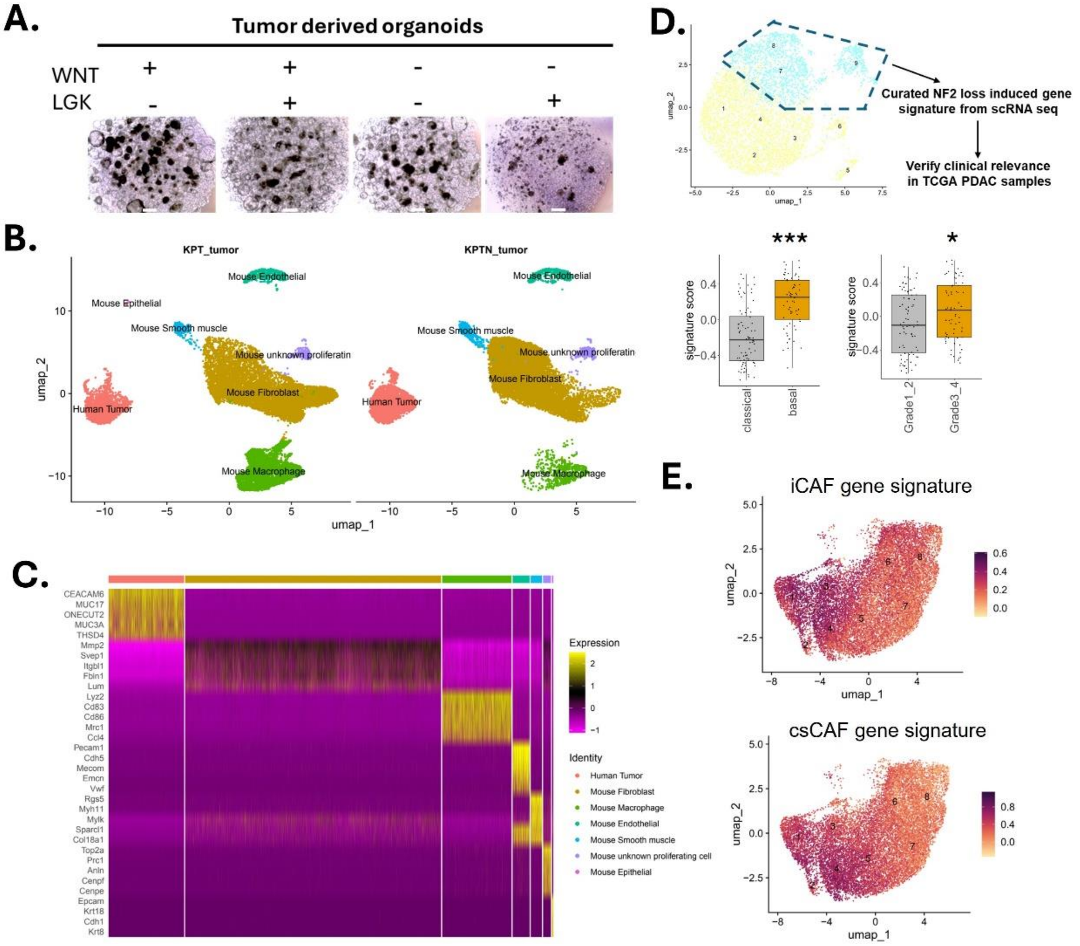


**Figure S3. A.** Representative images of tumor-derived organoids incubated with or without WNT3a supplement or LGK974 treatment (100 nM). Scale bar = 500 µm. **B.** Annotation of all cell populations present in KPT and KPTN-derived tumor tissues in single cell RNA seq analysis. **C.** Expression heatmap of top feature genes in each cell populations as shown in **B**. **D.** Expression of *NF2* induced malignancy-associated genes curated from single cell RNA seq data in TCGA PDAC samples. PDAC samples were grouped by subtype or grade. *, *** indicate p-value < 0.05 and 0.001, respectively. **E.** Expression of indicated gene modules in all the populations as shown in **B**.

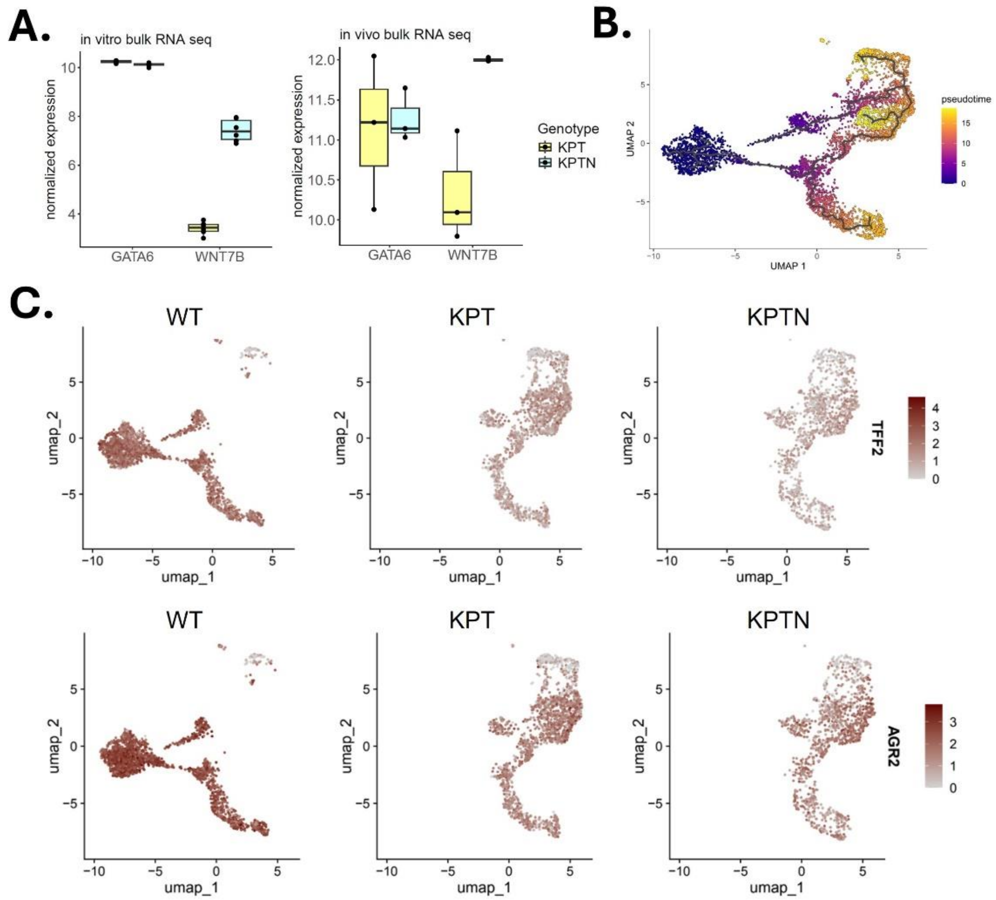


**Figure S4. A.** Log2 transformed normalized gene expression of *WNT7B* and *GATA6* in bulk RNA seq analysis of KPT and KPTN *in vitro* cultures (left, n = 6 each) and xenograft tumors (right, n = 3 each). **B.** Pseudo time trajectory analysis of all cell populations shown in Figure **4E**. **C.** Expression level of indicated genes in all the cells populations shown in Figure **4E**.

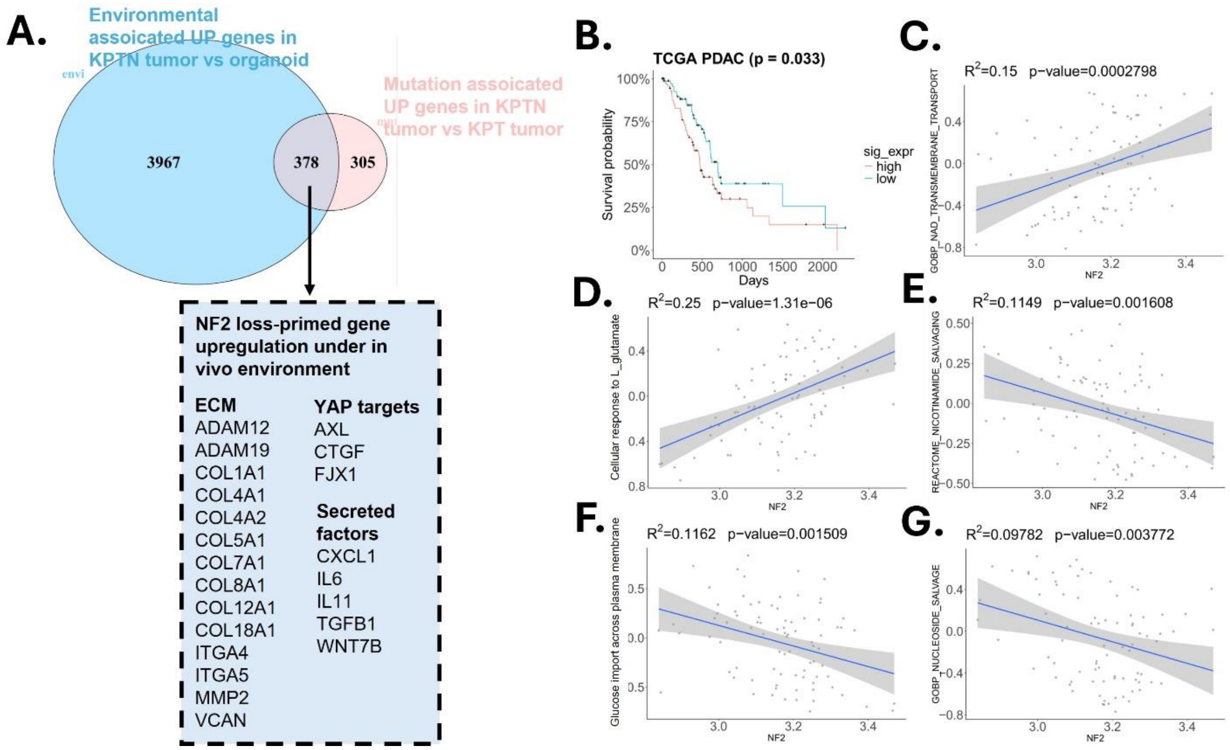


**Figure S5. A.** Comparison of environmentally and genetically induced upregulated genes in KPTN tumors. Environmentally induced upregulated genes were curated from the comparison of KPTN tumors vs organoids (shown in Figure **5A**). *NF2* loss induced upregulated genes was curated from the comparison of KPTN tumors vs KPT tumors (shown in Figure **3A**). **B.** KM plot of TCGA PDAC patient survival separated by median expression score of starvation associated genes as described in Figure **5A-B**. **C-G.** Correlation of indicated pathways with *NF2* expression level in TCGA PDAC samples.

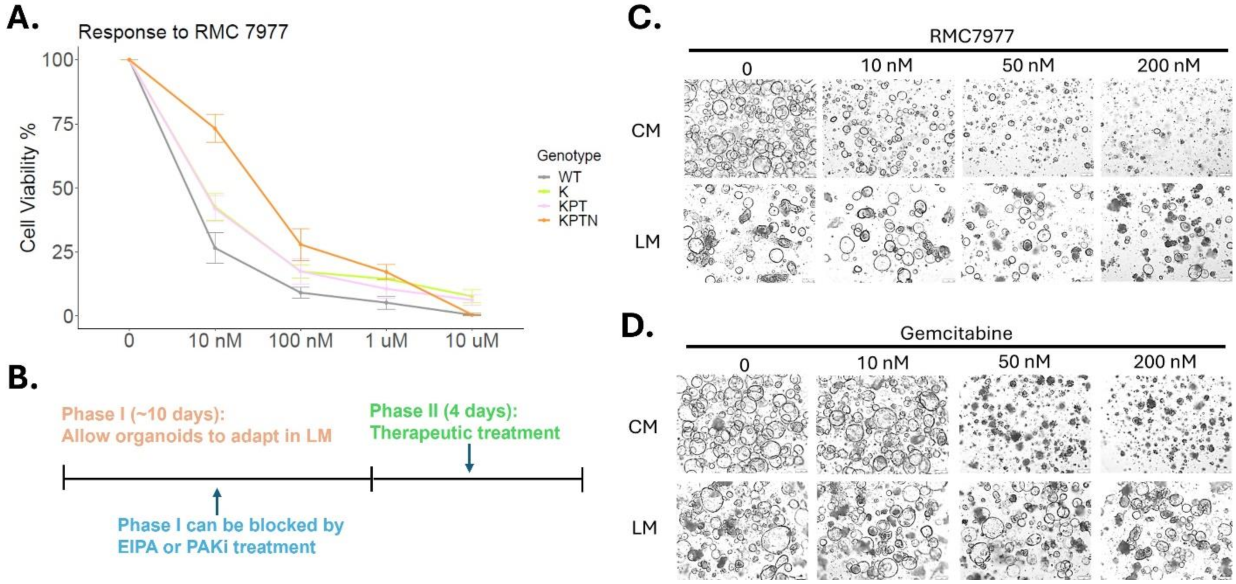


**Figure S6. A.** Response to RMC 7977 from acinar 3D cultures harboring different genetic backgrounds. **B.** Schematic illustration of experimental workflow for **Figure 6A-F**. **C.** Representative images of KPTN organoids cultured in CM or LM and treated with RMC 7977 at indicated doses or vehicle control. Scale bar = 250 µm. **D.** Representative images of KPTN organoids cultured in CM or LM and treated with Gemcitabine at indicated doses or vehicle control. Scale bar = 250 µm.

### Supplementary Table List

**Supplementary Table 1.** Organ donor information of the pancreatic tissues used in this study.

**Supplementary Table 2.** sgRNA sequences of the CRISPR knockout library.

**Supplementary Table 3.** Enrichment analysis summary at gene level for the in vivo and in vitro CRISPR screen.

**Supplementary Table 4.** List of DEGs from bulk RNA seq analysis described in Figure 3A, 4A and 5A.

**Supplementary Table 5.** Top 20 marker genes for each cell population from scRNA seq analysis described in Figure 3E, 3I and 4E.

**Supplementary Table 6.** Other curated gene signatures from the present or previous studies.
